# Regulatory plasticity of the rdar biofilm morphotype in clinical uropathogenic *Escherichia coli* and its modulation by ciprofloxacin

**DOI:** 10.64898/2026.04.22.719398

**Authors:** Fizza Mushtaq, Irfan Ali, Hira Sarfaraz, Rehmat Younas, Anju Bala, Bernt Eric Uhlin, Ute Römling, Irfan Ahmad

## Abstract

**Background:** Red, dry and rough (rdar) biofilm formation in *Escherichia coli* is characterized by the coordinated production of extracellular cellulose and amyloid curli fimbriae. Although rdar biofilm formation has been extensively characterized in laboratory strains, its clinical relevance and implications for antimicrobial treatment remain poorly understood.

**Methods:** We screened 146 consecutive *E. coli* isolates from patients with urinary tract infections in Lahore, Pakistan (2017–2019) for Congo red morphotypes at 28 °C and 37 °C. Antimicrobial susceptibility was determined by disc diffusion and compared between morphotypes using Fisher’s exact test. Fourteen isolates were whole-genome sequenced, the *csgD*–*csgBA* intergenic region was aligned and csgD expression was quantified by qRT-PCR. The effect of 10 µg/mL ciprofloxacin on rdar morphotype expression, matrix ultrastructure and biofilm formation was examined in ciprofloxacin-resistant isolates (MIC ≥ 128 µg/mL).

**Results:** Rdar-like colony morphotypes were expressed by 36.3% (53/146) of isolates at 28 °C and 29.5% (43/146) at 37 °C. Nucleotide polymorphisms in the *csgD*–*csgBA* intergenic region clustered within annotated OmpR, IHF and CsgD binding regions and were associated with, but did not individually predict, the mode of rdar morphotype expression. Isolates carrying the rdar morphotype were distributed across diverse *E. coli* phylogroups and sequence types. Resistance to aztreonam, cefuroxime and cephradine was more frequent among smooth and white isolates, whereas resistance to nitrofurantoin was more frequent among isolates rdar-positive at 28 °C (45.2% versus 20.2%, *p* = 0.0097); fluoroquinolone resistance did not differ between morphotypes. Subinhibitory ciprofloxacin (10 µg/mL) suppressed rdar biofilm formation, disrupted extracellular matrix architecture and abolished surface colonization on glass, in ciprofloxacin-resistant isolates.

**Conclusions:** Our findings establish the rdar morphotype as a clinically relevant biofilm phenotype in uropathogenic *E. coli* and show that subinhibitory ciprofloxacin suppresses rdar biofilm formation in ciprofloxacin-resistant isolates. These results underscore the importance of considering antibiotic-mediated modulation of biofilm behavior when interpreting treatment responses and designing strategies to combat persistent urinary tract infections.

## Background

Biofilms are structured communities of microorganisms embedded within a self-produced extracellular matrix comprised of exopolysaccharides, proteins, and nucleic acids. This multicellular lifestyle confers remarkable tolerance to environmental stresses, antibiotics, and host defenses, and is a major contributor to chronic and persistent infections in clinical settings [1–3].

Among the well-studied biofilm phenotypes in *Enterobacteriaceae* is the rdar (red, dry and rough) colony morphology, which is characterized by the production of adhesive extracellular matrix components including amyloid curli fimbriae and cellulose [4]. These biofilm matrix elements differentially facilitate surface adhesion, intercellular cohesion and pathogen-host interactions, hallmarks of mature biofilms that enhance resilience against external threats. The rdar morphotype is a widely occurring robust multicellular biofilm conserved among *Escherichia coli* and *Salmonella* species [5–7]. Rdar biofilm components and regulation are relevant in the colonization of diverse ecological niches such as plant tissue, tumor and immune cells, and play a significant role in long-term survival under harsh conditions such as desiccation [8–11].

The orphan transcriptional regulator CsgD is the central regulatory hub driving rdar morphotype expression in *Enterobacteriaceae* [12, 13]. Unphosphorylated CsgD directly activates expression of the curli subunit operon (*csgBA*C) and indirectly stimulates cellulose production by upregulating the diguanylate cyclase AdrA, which synthesizes the secondary messenger cyclic di-GMP (c-di-GMP). The c-di-GMP produced by AdrA post-translationally activates cellulose biosynthesis, while elevated c-di-GMP levels also act on downstream effectors to enhance *csgD* expression and matrix production and to repress motility, thereby reinforcing a sessile, biofilm-associated lifestyle [14, 15].

Regulation of *csgD* expression is highly complex. It involves a network of multiple global transcription factors, small regulators, and environmental cues, including temperature, nutrient status, and stress conditions [12, 13]. For instance, *csgD* expression is controlled by at least ten distinct regulatory inputs and is influenced by both transcriptional and post-transcriptional mechanisms, such as small RNAs and RNA-binding proteins (e.g., CsrA), as well as c-di-GMP signaling modules [16, 17]. These regulatory layers allow bacteria to finely tune biofilm formation in response to fluctuating conditions [18].

Natural isolates are heterogeneous with respect to their ability to express the rdar morphotype [5]. For instance, clinical isolates can express a temperature-dependent rdar morphotype, a semi-constitutive rdar morphotype, or no rdar morphotype at all, the last appearing as a smooth and white phenotype [19]. Modulation mediated through cyclic di-GMP signaling and mutations in the *csgD* promoter are two known regulatory determinants responsible for this variability in rdar morphotype expression [19–21]. Furthermore, natural isolates can concomitantly express both extracellular matrix components or only cellulose or curli fimbriae.

Despite intensive molecular characterization of the rdar biofilm regulatory network in laboratory and model strains, there remains a significant gap in our understanding of its relevance in clinical settings. Many studies of biofilm regulation, including those focused on *csgD* and c-di-GMP networks, have been carried out under controlled experimental conditions or using model isolates. In contrast, data on the prevalence, regulation, and functional consequences of rdar biofilm expression in clinical isolates are limited, particularly regarding how regulatory variation influences antimicrobial tolerance.

Although rdar biofilm formation has been linked to increased environmental persistence and transmission in non-clinical contexts, loss of the capacity to produce cellulose has been associated with a gain in virulence, suggesting a trade-off between environmental persistence and virulence [22]. Direct evidence of how variation in rdar regulatory networks affects clinical presentation or how rdar biofilms respond to antimicrobial treatment in the human host is, however, sparse. This gap underscores the need for studies that bridge mechanistic biofilm biology and clinical microbiology.

This manuscript aims to address these knowledge gaps by combining the investigation of prevalence of the rdar phenotype in clinically relevant strains with mechanistic insights along with investigation of the response of rdar morphotype expressing *E. coli* to antibiotics using ciprofloxacin as a relevant antibiotic.

## Methods

### Collection of *Escherichia coli* clinical isolates

Consecutive, well-isolated *E. coli* strains were collected from the urine of patients with urinary tract infection attending the Unique Diagnostic Laboratory, Lahore, Pakistan, between 2017 and 2019.

The study included a total of 146 UTI patients. Ethical approval and informed consent are described in the Ethics statement. The identity of all isolates was confirmed by biochemical methods. For that, a loop from the vial of preserved isolates was streaked on MacConkey agar. Plates were incubated at 37 °C overnight and the following day, the lactose fermenting isolates appeared as pink colonies on MacConkey agar which were processed for biochemical identification of *E. coli* as described [23]. *Escherichia coli* strains were subsequently stored in LB broth containing 15% glycerol at −80 °C. Selected isolates were further confirmed by whole genome sequence analysis.

### Antimicrobial susceptibility testing

A single colony was picked with a sterile loop from an overnight MacConkey plate and emulsified in sterile saline until no solid material from the colony remained visible. The turbidity of the suspension was adjusted to a 0.5 McFarland standard, corresponding to approximately 1–2 × 10⁸ CFU/mL, verified by measuring the optical density at 600 nm. A sterile cotton swab was dipped into the suspension and the excess fluid was removed by pressing the swab against the inside wall of the tube.

The swab was then used to streak a Mueller–Hinton agar plate to produce a lawn of growth, and the plates were left to dry for about 5 minutes. Antibiotic discs were applied to the agar surface at the correct spacing using flame-sterilized forceps. The panel comprised imipenem (10 µg), meropenem (10 µg), piperacillin–tazobactam (100/10 µg), ampicillin (10 µg), amoxicillin–clavulanic acid (20/10 µg), cephradine (30 µg), cefuroxime (30 µg), ceftriaxone (30 µg), cefaclor (30 µg), cefoperazone (75 µg), aztreonam (30 µg), amikacin (30 µg), gentamicin (10 µg), tetracycline (30 µg), ciprofloxacin (5 µg), norfloxacin (10 µg), trimethoprim–sulfamethoxazole (1.25/23.75 µg) and nitrofurantoin (300 µg). Not every isolate was tested against every agent. The plates were incubated for 16–18 hours at 37 °C.

The diameter of the zone of inhibition around each disc was measured with a ruler and interpreted as resistant, intermediate or susceptible according to the breakpoints of the Clinical and Laboratory Standards Institute (CLSI) for each agent. *E. coli* ATCC 25922 was included as a quality-control strain with each batch of tests.

### Determination of minimum inhibitory concentration

Minimum inhibitory concentrations of ciprofloxacin were determined by broth microdilution in cation-adjusted Mueller–Hinton broth according to CLSI methodology, using two-fold dilutions from 0.06 to 128 µg/mL and an inoculum of approximately 5 × 10⁵ CFU/mL. Plates were read after 16–20 hours at 37 °C and the MIC was taken as the lowest concentration with no visible growth. *E. coli* ATCC 25922 was included as a control.

### Detection of extended-spectrum β-lactamase production

ESBL production was detected phenotypically by the combination disc method, comparing the inhibition zone of ceftazidime and cefotaxime discs alone with that of the same discs combined with clavulanic acid. An increase of ≥ 5 mm in zone diameter in the presence of clavulanic acid was recorded as ESBL positive, following CLSI criteria.

### Detection of virulence determinants

The virulence genes *fimH*, *pap*, *sfa*, *afa*, K1 and K5 were detected by PCR on boiled-cell lysates using previously published primers and cycling conditions. Amplicons were resolved on agarose gels and sized against a molecular weight marker, with positive and negative control strains included in each run.

### Rdar morphotype formation assay

*E. coli* strains were tested for their capacity to express the rdar colony morphotype on salt-free LB agar supplemented with Congo red (40 μg ml−1) and Coomassie brilliant blue G (20 μg ml−1). A loop of bacterial growth from each strain was spotted onto two Congo red agar plates, incubated at 28 °C and 37 °C respectively, and colony morphology was recorded after 24, 48 and 72 hours. Colonies were scored into four classes [4, 24, 25]: red, dry and rough (rdar), indicating the expression of both curli and cellulose; pink, dry and rough (pdar), indicating cellulose in the absence of curli; brown, dry and rough (bdar), indicating curli in the absence of cellulose; and smooth and white (saw), indicating the expression of neither. Isolates expressing the rdar morphotype at both 28 °C and 37 °C are referred to as semi-constitutive and those expressing it only at 28 °C as temperature-dependent. Isolates forming the full rdar morphotype at 28 °C are referred to as rdar-positive, and the three matrix-expressing classes together (rdar, pdar and bdar) as rdar-like.

### Biofilm formation assay

Biofilm formation was also quantified in sterilized 96 well microtiter plates (Nunc™ Cell culture treated, Thermo scientific) as described previously [26, 27]. A suspension of bacterial cells grown overnight on LB agar plates was prepared in PBS with an OD600 = 1. From this suspension, twenty microliters were added to each well of the 96 well plates containing 180 microliters of salt-free LB broth, with or without 10 µg/mL ciprofloxacin as indicated. Plates were incubated in a moist chamber at 28 °C or 37 °C for 24 hours; the chamber was humidified to avoid evaporation from the peripheral wells. Subsequently, the liquid was discarded and the plates washed gently with water. Bacterial cells attached to walls, or at the base, of wells in the form of biofilm were stained with 1% crystal violet for 30 minutes. Excessive crystal violet was removed by washing three times with water. Biofilms stained with crystal violet were dissolved in 5% acetic acid solution. The concentration of dissolved crystal violet measured as optical density at 595 nm correlates with the abundance of biofilm. Data were subjected to statistical analysis using GraphPad Prism software.

### Whole-genome sequencing

Fourteen isolates were selected for whole-genome sequencing to span the range of Congo red phenotypes observed. Sequencing was performed using the MiSeq Desktop Sequencer and MiSeq Reagent Kit v3 (Illumina, San Diego, CA, USA) as described [28]. DNA preparation, library construction, and genome sequencing were done according to the manufacturers’ instructions. Sequence data were assembled and analyzed using the CLC genomics workbench (v7.0.4; CLC bio, Aarhus, Denmark). The web server Reference sequence Alignment-based Phylogeny builder (REALPHY) was used to construct a phylogenetic tree based on multiple alignments of the genomic sequence data [29].

The MLST web-based search engine, hosted by the Center for Genomic Epidemiology in Denmark (http://www.genomicepidemiology.org/), was used to assign the isolates into sequence types (STs) [30]. The occurrence of acquired antimicrobial resistance genes was detected using the ResFinder service, also hosted by the Center for Genomic Epidemiology in Denmark [31]. The occurrence of resistance and virulence genes was verified, and genetic surroundings were annotated based on the yields of nucleotide similarities obtained using the Basic Local Alignment Search Tool (http://blast.ncbi.nlm.nih.gov/Blast.cgi) against the “Nucleotide collection (nr/nt)” and/or “Whole-genome shotgun contigs (wgs)” databases [32].

### Scanning electron microscopy

Sample preparation and scanning electron microscopy were performed as described [27]. Briefly, five microliters of a bacterial suspension in PBS (OD600 of 1) from an overnight agar plate culture were spotted onto salt-free LB agar supplemented with Congo red and Coomassie brilliant blue G. Pieces of agar carrying bacterial colonies were excised and fixed overnight at 4 °C in 2.5% glutaraldehyde in 0.1 M sodium cacodylate, dehydrated through a graded ethanol series, critical-point dried and coated with 5 nm gold/palladium. Cell morphology was examined with a field-emission scanning electron microscope (Carl Zeiss Merlin FESEM) using secondary electron detectors at an accelerating voltage of 4 kV and a probe current of 50–100 pA.

### qRT-PCR

RNA was purified from bacteria grown on salt-free LB plates at 28 °C using the Total RNA Isolation kit (Qiagen) according to the manufacturer’s protocol. After determination of RNA concentration with a NanoDrop ND-1000 UV-Vis spectrophotometer, 1 μg of RNA was reverse transcribed in a 20 μl reaction using a cDNA Reverse Transcription Kit (Sigma Aldrich). Twenty nanograms of template were used for real-time PCR with Power SYBR Green PCR Master Mix (Sigma Aldrich). Cycling was performed on an ABI 7500 Real Time PCR System. Individual gene expression profiles were normalized against the 16S rRNA gene as endogenous control. Values are means of at least three independent experiments with two technical replicates and error bars represent standard deviations; statistical evaluation is described under Statistical Analysis.

### Biofilm formation on glass surfaces

For analyses of colonization of *E. coli* on glass surfaces, the assay was performed on sterilized 18-well chamber glass slides (Ibidi GmbH) as described previously with few modifications [33]. Bacterial cells grown overnight on LB agar plates were suspended in PBS to an OD600 = 1. From this suspension, 20 µl of bacterial suspension were added to each well, containing 180 µl of salt-free LB broth supplemented with 10 µg/mL ciprofloxacin or without ciprofloxacin. Chamber slides were incubated in a moist chamber at 28 °C. Subsequently, the liquid content was removed, and the plates were washed three times with PBS. Bacterial cells attached to the glass surface in the form of biofilm were visualized using phase contrast microscopy and subsequently stained with 1% crystal violet for 30 min. Excess crystal violet was removed by washing three times with water. Cells stained with crystal violet were dissolved in 1 ml of 5% acetic acid solution. The intensity of the crystal violet color represents the abundance of biofilm estimated as optical density at 595 nm. Planktonic growth curves were recorded in LB with and without 10 µg/mL ciprofloxacin at 37 °C using the temperature-control mode of a Spark multimode plate reader (Tecan), with OD600 recorded at 60 min intervals for 12 hours in triplicate. Data were subjected to statistical analysis using GraphPad Prism software.

### Statistical analysis

Differences in resistance rates between morphotype groups were assessed with Fisher’s exact test, which was preferred to the chi-square test because several cells contained small expected counts. Differences in *csgD* mRNA level between phenotypic classes were assessed by one-way ANOVA followed by Tukey’s multiple comparisons test. Biofilm quantification in microtiter plates (Figure 5C) and on glass surfaces (Figure 6C) were compared using a two-tailed unpaired t-test with Welch’s correction. Analyses were performed in GraphPad Prism and in Python (SciPy). A *p* value < 0.05 was considered statistically significant.

## Results

### Simultaneous expression of cellulose and curli fimbriae is required for intact rdar biofilm architecture

To analyze the rdar morphotype at the single-cell level, rdar biofilms formed by *E. coli* No. 12 [34], a well-characterized uropathogenic rdar-biofilm forming clinical isolate, and its mutant derivatives deficient in cellulose or amyloid curli fimbriae production were visualized using scanning electron microscopy (SEM). SEM imaging revealed that biofilm matrix components enveloped individual bacterial cells in a continuous, honeycomb-like sheet, when both cellulose and curli fimbriae were simultaneously expressed (Figure 1). In contrast, the absence of either cellulose or curli fimbriae appeared to alter matrix integrity, resulting in disruption of the cohesive covering and dispersal of extracellular matrix material across the bacterial population (Figure 1). Because sample preparation for SEM involves fixation, dehydration and critical-point drying, which can collapse or redistribute the extracellular matrix, these differences are interpreted qualitatively. This finding is consistent with earlier observations that rdar morphotype extracellular matrix components are deposited around individual cells within biofilms [35]. This phenomenon is distinct from *Acinetobacter baumannii* or *Pseudomonas aeruginosa* biofilms where Csu pili from the neighboring cells establish antiparallel interactions to form 3D mature biofilms [36].

**Figure 1.**
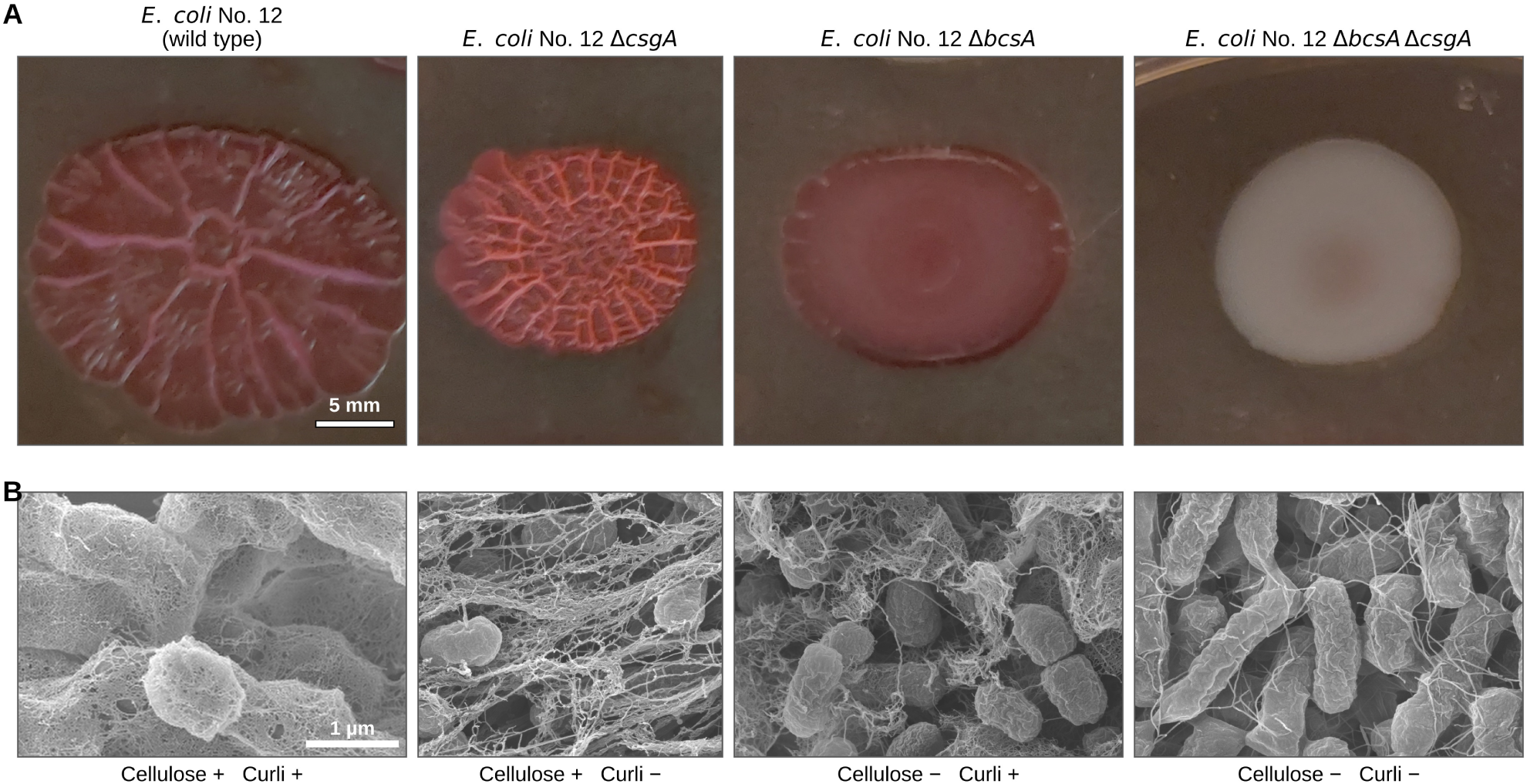
Expression of rdar biofilm components in *Escherichia coli* No. 12. (A) Representative images of rdar biofilm formation by *E. coli* No. 12 and its mutant derivatives after 72 h of growth on Congo red agar at 28 °C. All four panels are crops of a single plate recorded in one photograph; one uniform flat-field correction was applied to the whole image before cropping and each panel is reproduced at the same magnification and at the native pixel aspect ratio. Scale bar, 5 mm, applies to all panels. (B) Scanning electron microscopy (SEM) images of *E. coli* No. 12 and mutant strains grown on Congo red agar for 72 h at 28 °C, illustrating ultrastructural differences in extracellular matrix organization. The wild-type micrograph was resampled to the scale of the mutant panels; the original is shown in Additional file 4: Figure S1. Scale bar, 1 µm, applies to all panels.

### Frequency of rdar morphotype formation among clinical *E. coli* urinary isolates

Next, we assessed the frequency of rdar biofilm formation among clinical *E. coli* isolates recovered from urinary tract infection (UTI) patients to evaluate the clinical relevance of this phenotype. A total of 146 consecutive *E. coli* isolates collected from UTI patients attending outpatient clinics in Lahore, Pakistan, between 2017 and 2019 were analyzed. Per-isolate morphotype, susceptibility and virulence data are given in the Additional file 1: Table S1.

Assessment of Congo red binding on agar plates revealed that 36.3% (53/146) of isolates expressed a rdar-like biofilm morphotype at 28 °C. Of these, 22.6% (33/146) formed a pronounced complete red, dry and rough (rdar) morphotype, 4.1% (6/146) a pink, dry and rough (pdar) morphotype and 9.6% (14/146) a brown, dry and rough (bdar) morphotype; the remaining 63.7% (93/146) formed smooth and white colonies (Figure 2A).

**Figure 2.**
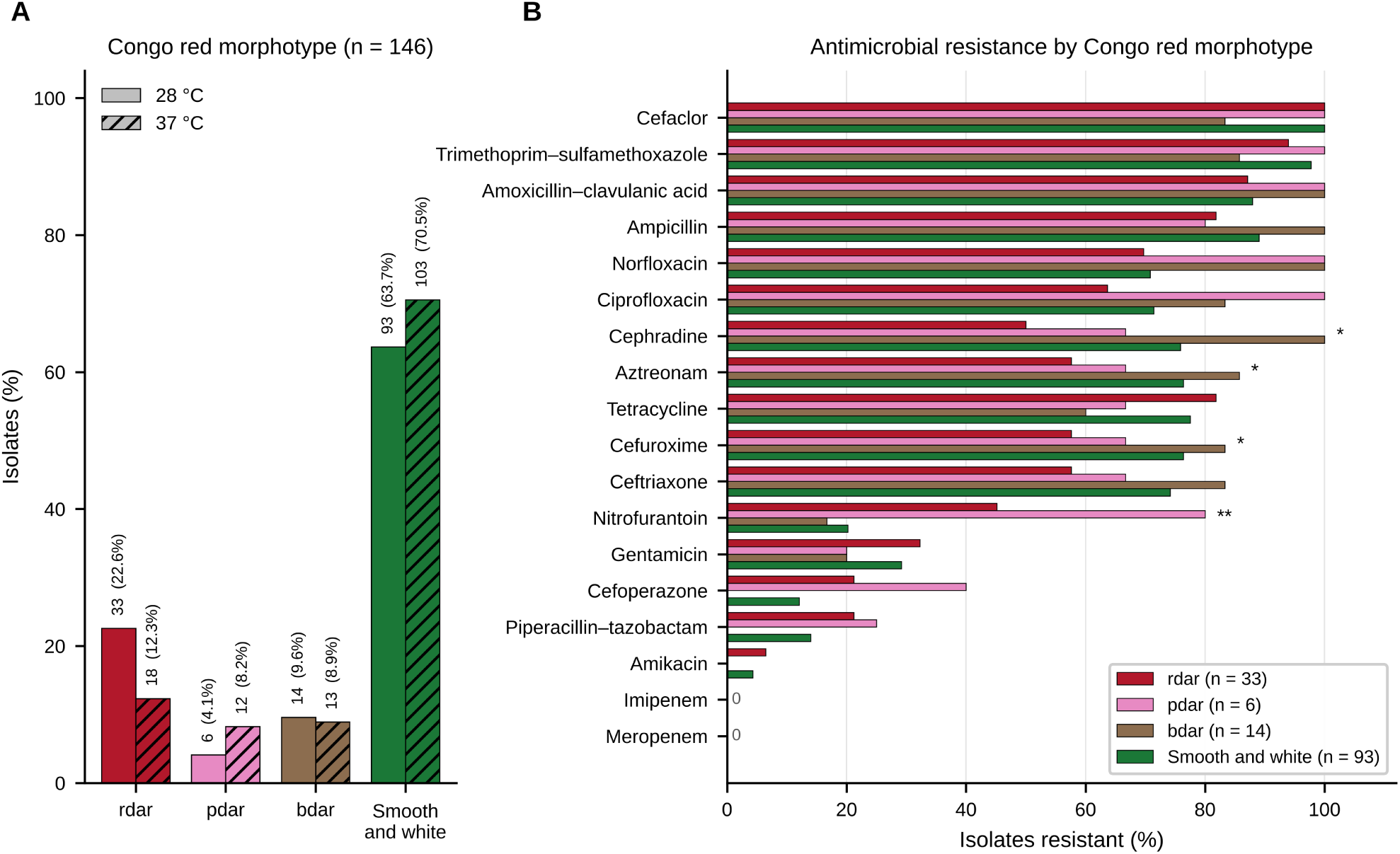
Distribution of Congo red morphotypes and their relationship to antimicrobial resistance among clinical *Escherichia coli* urinary isolates. (A) Proportion of isolates expressing each Congo red morphotype after incubation at 28 °C and at 37 °C; the number of isolates is given before the percentage, *n* = 146 at each temperature. (B) Resistance rate for each antimicrobial agent, stratified by Congo red morphotype, among the 146 isolates subjected to antimicrobial susceptibility testing. Group sizes are given in the key, and agents for which no isolate was resistant are marked 0. Asterisks denote a significant difference in resistance between rdar-positive and smooth and white isolates by Fisher’s exact test: * *p* < 0.05, ** *p* < 0.01. In (B), each percentage is calculated over the isolates of that morphotype group tested against the agent; the numbers tested are given in Table 1.

**Table 1.** Antimicrobial susceptibility of clinical *Escherichia coli* urinary isolates in relation to Congo red morphotype. Values are the number of resistant / intermediate / susceptible isolates, with the percentage resistant in parentheses. Denominators differ between agents because not every isolate was tested against every agent; where fewer than the full group was tested the number tested is shown in brackets. Susceptibility testing was performed at 37 °C; Congo red morphotype was scored at 28 °C.

| **Antimicrobial agent** | **rdar (*n* = 33)** | **pdar (*n* = 6)** | **bdar (*n* = 14)** | **saw (*n* = 93)** | **All isolates (*n* = 146)** |
|---|---|---|---|---|---|
| Imipenem | 0 / 0 / 31 (0.0%) [31] | 0 / 0 / 5 (0.0%) [5] | 0 / 0 / 14 (0.0%) | 0 / 0 / 93 (0.0%) | **0 / 0 / 143 (0.0%)** |
| Meropenem | 0 / 0 / 33 (0.0%) | 0 / 0 / 5 (0.0%) [5] | 0 / 0 / 14 (0.0%) | 0 / 0 / 93 (0.0%) | **0 / 0 / 145 (0.0%)** |
| Piperacillin–tazobactam | 7 / 0 / 26 (21.2%) | 1 / 1 / 2 (25.0%) [4] | 0 / 0 / 14 (0.0%) | 13 / 4 / 76 (14.0%) | **21 / 5 / 118 (14.6%)** |
| Ampicillin | 27 / 0 / 6 (81.8%) | 4 / 0 / 1 (80.0%) [5] | 14 / 0 / 0 (100.0%) | 81 / 0 / 10 (89.0%) [91] | **126 / 0 / 17 (88.1%)** |
| Amoxicillin–clavulanic acid | 27 / 0 / 4 (87.1%) [31] | 6 / 0 / 0 (100.0%) | 14 / 0 / 0 (100.0%) | 73 / 2 / 8 (88.0%) [83] | **120 / 2 / 12 (89.6%)** |
| Cephradine | 10 / 0 / 10 (50.0%) [20] | 2 / 0 / 1 (66.7%) [3] | 8 / 0 / 0 (100.0%) [8] | 44 / 0 / 14 (75.9%) [58] | **64 / 0 / 25 (71.9%)** |
| Cefuroxime | 19 / 0 / 14 (57.6%) | 4 / 0 / 2 (66.7%) | 10 / 0 / 2 (83.3%) [12] | 71 / 0 / 22 (76.3%) | **104 / 0 / 40 (72.2%)** |
| Ceftriaxone | 19 / 0 / 14 (57.6%) | 4 / 0 / 2 (66.7%) | 10 / 0 / 2 (83.3%) [12] | 69 / 0 / 24 (74.2%) | **102 / 0 / 42 (70.8%)** |
| Cefaclor | 31 / 0 / 0 (100.0%) [31] | 5 / 0 / 0 (100.0%) [5] | 10 / 0 / 2 (83.3%) [12] | 91 / 0 / 0 (100.0%) [91] | **137 / 0 / 2 (98.6%)** |
| Cefoperazone | 7 / 0 / 26 (21.2%) | 2 / 1 / 2 (40.0%) [5] | 0 / 0 / 14 (0.0%) | 11 / 4 / 76 (12.1%) [91] | **20 / 5 / 118 (14.0%)** |
| Aztreonam | 19 / 0 / 14 (57.6%) | 4 / 0 / 2 (66.7%) | 12 / 0 / 2 (85.7%) | 71 / 0 / 22 (76.3%) | **106 / 0 / 40 (72.6%)** |
| Amikacin | 2 / 0 / 29 (6.5%) [31] | 0 / 0 / 5 (0.0%) [5] | 0 / 0 / 14 (0.0%) | 4 / 2 / 87 (4.3%) | **6 / 2 / 135 (4.2%)** |
| Gentamicin | 10 / 0 / 21 (32.3%) [31] | 1 / 0 / 4 (20.0%) [5] | 2 / 0 / 8 (20.0%) [10] | 26 / 2 / 61 (29.2%) [89] | **39 / 2 / 94 (28.9%)** |
| Tetracycline | 27 / 0 / 6 (81.8%) | 4 / 0 / 2 (66.7%) | 6 / 2 / 2 (60.0%) [10] | 69 / 0 / 20 (77.5%) [89] | **106 / 2 / 30 (76.8%)** |
| Ciprofloxacin | 21 / 0 / 12 (63.6%) | 6 / 0 / 0 (100.0%) | 10 / 0 / 2 (83.3%) [12] | 65 / 0 / 26 (71.4%) [91] | **102 / 0 / 40 (71.8%)** |
| Norfloxacin | 23 / 0 / 10 (69.7%) | 6 / 0 / 0 (100.0%) | 10 / 0 / 0 (100.0%) [10] | 63 / 0 / 26 (70.8%) [89] | **102 / 0 / 36 (73.9%)** |
| Trimethoprim–sulfamethoxazole | 31 / 0 / 2 (93.9%) | 6 / 0 / 0 (100.0%) | 12 / 0 / 2 (85.7%) | 87 / 0 / 2 (97.8%) [89] | **136 / 0 / 6 (95.8%)** |
| Nitrofurantoin | 14 / 0 / 17 (45.2%) [31] | 4 / 0 / 1 (80.0%) [5] | 2 / 0 / 10 (16.7%) [12] | 18 / 0 / 71 (20.2%) [89] | **38 / 0 / 99 (27.7%)** |

rdar, red dry and rough; pdar, pink dry and rough; bdar, brown dry and rough; saw, smooth and white.
Cephradine was tested against a subset of isolates only.

At 37 °C, 29.5% (43/146) of isolates still bound Congo red. Among these, 18 isolates (12.3%) displayed the rdar morphotype, 12 isolates (8.2%) the pdar morphotype and 13 isolates (8.9%) the bdar morphotype, while 103 isolates (70.5%) formed smooth and white colonies (Figure 2A). Isolates that expressed a rdar morphotype at both temperatures are referred to below as semi-constitutive, and those expressing it only at 28 °C as temperature-dependent.

### Whole-genome sequencing–based phylogenetic analysis reveals that rdar morphotype– forming isolates are distributed across the *E. coli* phylogeny

To further investigate the genetic basis of rdar morphotype expression in relation to phylogeny, whole-genome sequencing was performed on fourteen clinical isolates selected to span the range of Congo red phenotypes observed. Whole-genome sequencing–based analysis of sequence types, as well as the distribution of antimicrobial resistance and virulence genes in these isolates are summarized in Additional file 1: Table S1. Comparative distributions of resistance and virulence genes relative to rdar morphotype are presented in Additional file 2: Table S2 and Additional file 3: Table S3, respectively.

A whole-genome sequence–based phylogenetic tree was constructed using reference strains with known phylogroups, including Fec101 (B1), MG1655 (A), B8638 (D), TOB1 (B2), B11870 (B2), Fec67 (B2), EcoR31 and *E. coli* No. 12 (B2). Rdar biofilm-forming isolates were distributed across all examined phylogroups and frequently clustered together with smooth and white isolates within the same phylogenetic groups (Figure 3). These findings indicate that rdar biofilm formation is not restricted to particular sequence types and occurs across the breadth of the *E. coli* phylogeny.

**Figure 3.**
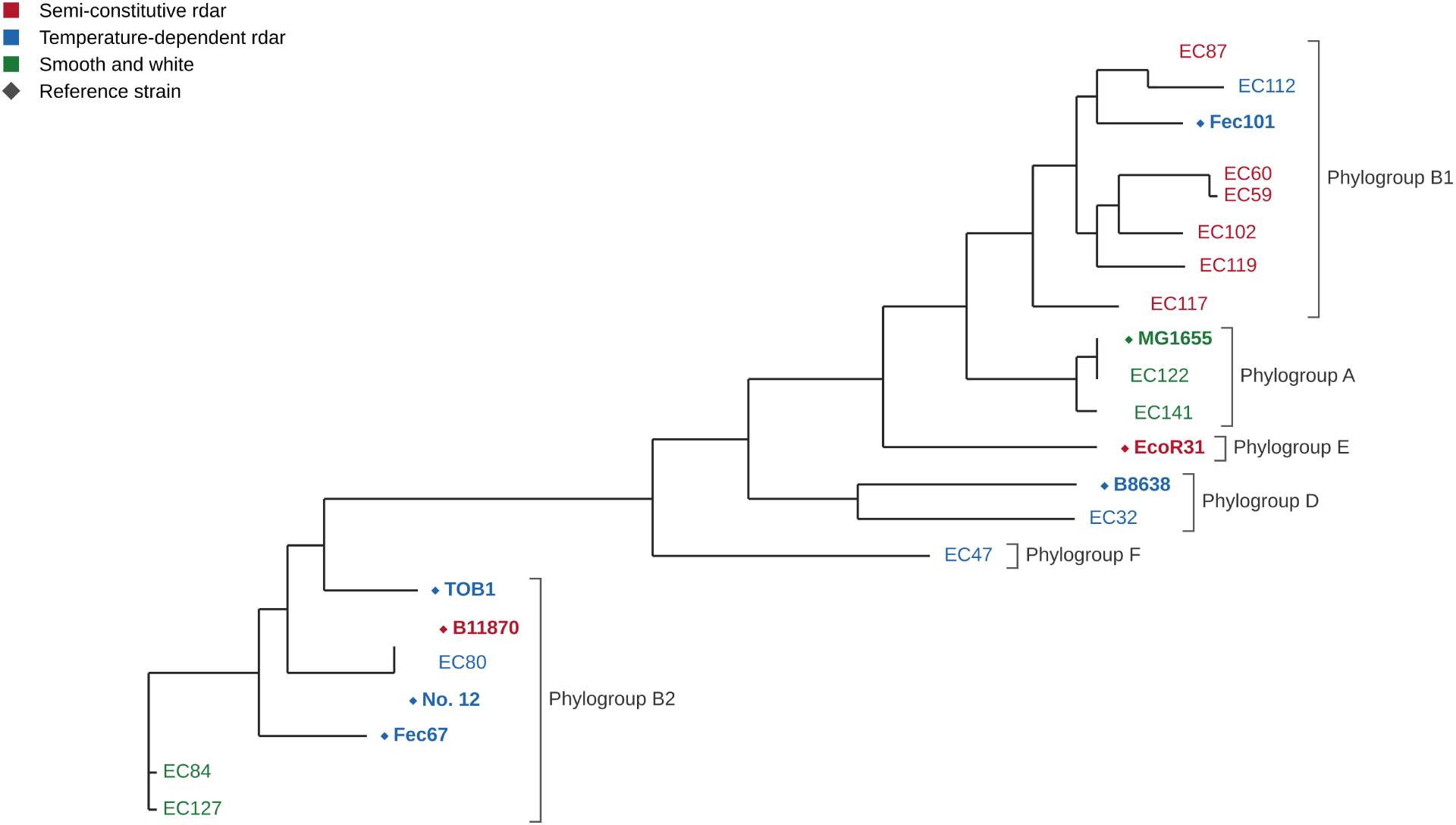
Phylogenetic analysis of representative *Escherichia coli* isolates. Whole-genome–based phylogenetic tree generated using REALPHY (https://realphy.unibas.ch/realphy/) from multiple genome sequence alignments. The genome of *E. coli* MG1655 (phylogroup A) was used as the reference sequence. Reference genomes used for phylogroup assignment include *E. coli* Fec101 (phylogroup B1), *E. coli* TOB1 (phylogroup B2), *E. coli* B11870 (phylogroup B2), *E. coli* Fec67 (phylogroup B2), *E. coli* No. 12 (phylogroup B2) and *E. coli* B8638 (phylogroup D); *E. coli* EcoR31 (phylogroup E) is included as a previously characterized semi-constitutive rdar strain. Tip labels are colored by Congo red phenotype: red, semi-constitutive rdar; blue, temperature-dependent rdar; green, smooth and white. Previously characterized reference strains are shown in bold and marked with a diamond, and brackets indicate phylogroup assignment; phylogroups of the study isolates were assigned from their position in the tree and their sequence type (Additional file 1: Table S1).

### Nucleotide polymorphism in the *csgD*–*csgBA* intergenic region is associated with the variability in rdar morphotype expression in clinical isolates

CsgD is a key regulator of rdar morphotype formation, and the intergenic region upstream of the *csgD* open reading frame serves as a binding platform for multiple global transcriptional regulators. Among those regulators, CpxR and H-NS act as repressors while OmpR, RstA and IHF act as activators [37]. We therefore compared the 754 bp *csgD*–*csgBA* intergenic sequences among the sequenced isolates. We also included the corresponding sequences of nine previously characterized reference strains retrieved from the database, which differ in their mode of rdar expression, giving 23 sequences in total. These strains are listed in Additional file 7: Table S4.

Multiple sequence alignment shows nucleotide polymorphism in the promoter region at the binding sites of OmpR, IHF and CsgD and at the ribosome binding site of the corresponding mRNA (Figure 4A–D).

**Figure 4.**
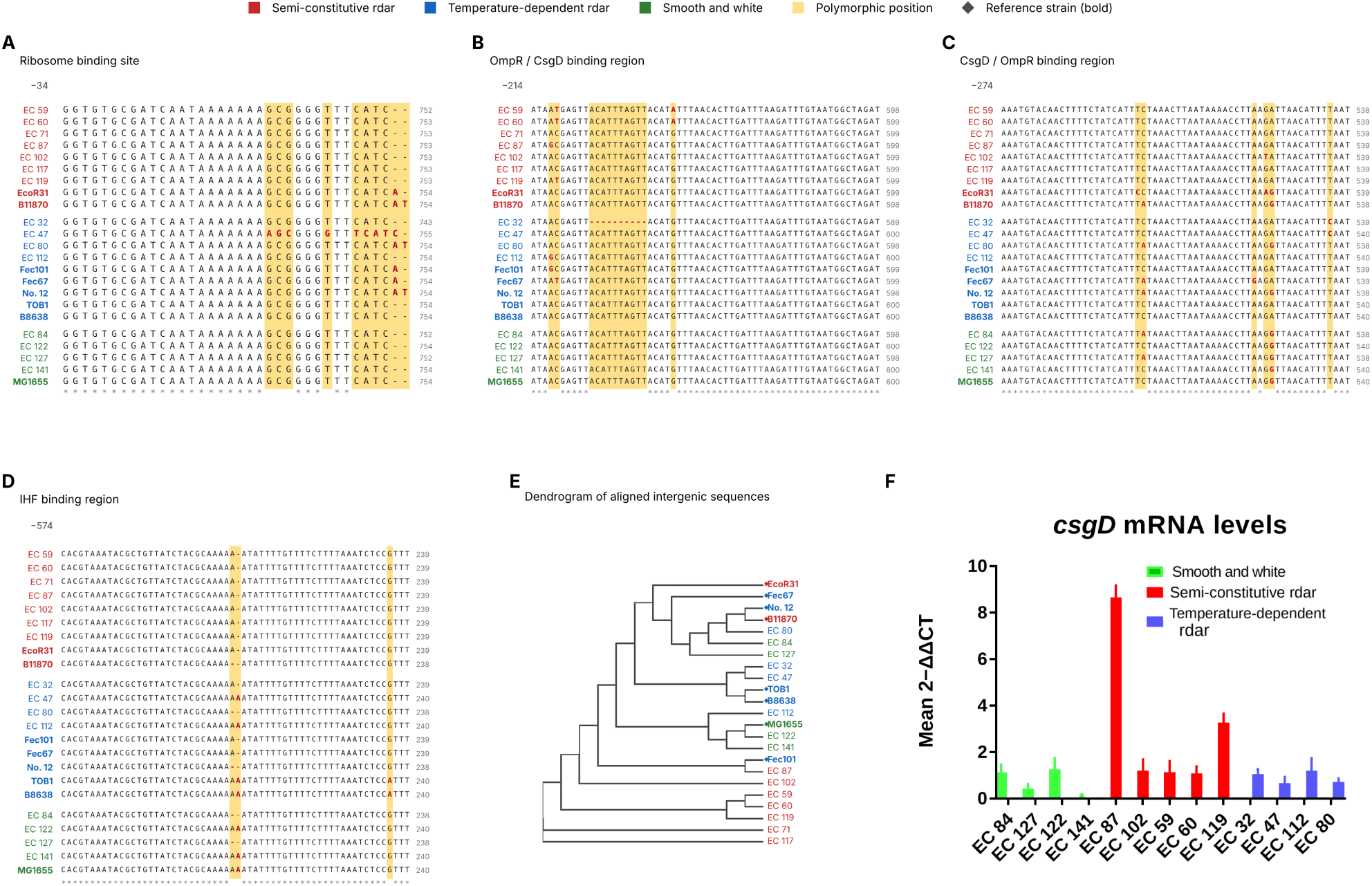
Sequence variation in the *csgD*–*csgBA* intergenic region and its association with rdar morphotype expression. (A–D) Regions of the multiple sequence alignment of the *csgD*–*csgBA* intergenic region spanning the ribosome binding site (A) and the annotated OmpR/CsgD (B), CsgD/OmpR (C) and IHF (D) binding regions, for all 23 aligned sequences (14 study isolates and 9 previously characterized reference strains from the database). Isolates are ordered by phenotypic class, polymorphic positions are shaded and bases differing from the majority are shown in bold red. Nucleotide positions indicated in panels A–D are relative to the translation start site of CsgD in *E. coli* MG1655, and the coordinate of the last base of each window is given at the right of each row. The complete alignment is provided as Additional file 5: Figure S2. Color code: green, smooth and white; red, semi-constitutive rdar; blue, temperature-dependent rdar. Previously characterized reference strains are shown in bold and marked with a diamond. (E) Phylogenetic tree of aligned *csgD*–*csgBA* intergenic sequences, constructed using *E. coli* MG1655 as the reference strain. (F) Relative *csgD* mRNA levels in bacteria grown on salt-free LB agar plates for 24 hours at 28 °C measured by quantitative RT-PCR. Values are means of at least three independent experiments with two technical replicates; error bars show standard deviations. Differences between phenotypic classes were assessed by one-way ANOVA followed by Tukey’s multiple comparisons test.

Dendrogram of aligned sequences indicates a clustering pattern of sequence types from semi-constitutive rdar morphotype isolates analyzed in this study. However, the sequences from the semi-constitutive rdar morphotype forming reference strains EcoR31 and B11870 did not fall in this cluster (Figure 4E). The polymorphisms are confined to the regulator binding regions, and sequences from the semi-constitutive rdar forming isolates of this collection tend to group together; we therefore report an association between nucleotide polymorphism in the *csgD*– *csgBA* intergenic region and the variability in rdar morphotype expression, while noting that no individual polymorphism segregates with phenotype across the full set of strains examined.

Further on, *csgD* expression in these isolates was assessed by measuring *csgD* mRNA levels by quantitative RT-PCR (Figure 4F). There was no significant difference in *csgD* mRNA levels between temperature-dependent rdar-positive isolates and smooth and white colony-forming isolates. However, *csgD* mRNA levels were significantly higher in some of the semi-constitutive rdar biofilm isolates, suggesting that more than one route to rdar biofilm formation operates in *E. coli*, as reported previously [19, 20].

### Antimicrobial susceptibility patterns in relation to rdar biofilm formation

Antimicrobial susceptibility testing was performed at 37 °C on all 146 isolates and revealed heterogeneous resistance profiles across the collection (Table 1, Figure 2B). No isolate was resistant to imipenem or meropenem. Resistance rates were 88.1% (126/143) to ampicillin, 89.6% (120/134) to amoxicillin–clavulanic acid, 14.6% (21/144) to piperacillin–tazobactam, 71.9% (64/89) to cephradine, 98.6% (137/139) to cefaclor, 72.2% (104/144) to cefuroxime, 70.8% (102/144) to ceftriaxone, 14.0% (20/143) to cefoperazone, 72.6% (106/146) to aztreonam, 4.2% (6/143) to amikacin, 28.9% (39/135) to gentamicin, 76.8% (106/138) to tetracycline, 71.8% (102/142) to ciprofloxacin, 73.9% (102/138) to norfloxacin, 95.8% (136/142) to trimethoprim–sulfamethoxazole and 27.7% (38/137) to nitrofurantoin. Because not every isolate was tested against every agent, the denominator for each agent is the number of isolates tested against it. All percentages in the text, in Table 1 and in Figure 2B are calculated on these per-agent denominators, and the number tested in each morphotype group is given in Table 1.

Comparing isolates by Congo red morphotype, resistance differed significantly between rdar-positive (rdar morphotype, *n* = 33) and smooth and white (*n* = 93) isolates for four of the agents tested (Fisher’s exact test). Resistance was significantly higher among smooth and white isolates for aztreonam (57.6% versus 76.3%, *p* = 0.047), cefuroxime (57.6% versus 76.3%, *p* = 0.047) and cephradine (50.0% versus 75.9%, *p* = 0.048), whereas resistance to nitrofurantoin was significantly higher among rdar-positive isolates (45.2% versus 20.2%, *p* = 0.0097). Nitrofurantoin resistance was also more frequent among the cellulose-only pdar isolates (80.0%, 4/5, versus 20.2%; *p* = 0.010), and considering all cellulose-producing isolates together (rdar and pdar), 50.0% (18/36) were resistant compared with 20.2% (18/89) of smooth and white isolates (*p* = 0.0019). Resistance among the curli-only bdar isolates (16.7%, 2/12) was similar to that of smooth and white isolates. Differences for the remaining agents did not reach statistical significance, including ciprofloxacin (63.6% versus 71.4%, *p* = 0.51) and norfloxacin (69.7% versus 70.8%, *p* = 1.00). All six pdar isolates were resistant to both ciprofloxacin and norfloxacin, although with this group size the difference from the remaining isolates was not significant (100% versus 70.6%, *p* = 0.18).

Virulence determinants and extended-spectrum β-lactamase (ESBL) production were also assessed in relation to Congo red morphotype (Additional file 6: Figure S3). The type 1 fimbrial adhesin encoding *fimH* was present in the great majority of isolates of every morphotype group (87–100%), whereas indicator genes for P- and S-fimbriae, the afimbrial adhesin and K1 and K5 capsules, *pap*, *sfa*, *afa*, K1 and K5, respectively, were infrequent. ESBL production was detected in 27.3% (9/33) of rdar, 16.7% (1/6) of pdar, 42.9% (6/14) of bdar and 48.4% (45/93) of smooth and white isolates.

### Ciprofloxacin suppresses rdar biofilm formation in ciprofloxacin-resistant isolates

Ciprofloxacin is one of the most commonly prescribed antibiotics for the treatment of UTIs, and in our collection every pdar isolate — a morphotype characterized by the production of cellulose in the absence of curli — was resistant to both ciprofloxacin and norfloxacin (Table 1). We therefore examined the effect of exogenous ciprofloxacin on rdar expression in three ciprofloxacin-resistant isolates (*E. coli* EC47, *E. coli* EC112 and *E. coli* EC119), each with a minimum inhibitory concentration (MIC) ≥ 128 µg/mL. The resistance-associated mutations in these isolates are summarized in Table 2.

**Table 2.** Mutations in the quinolone resistance-determining regions of *gyrA*, *gyrB*, *parC* and *parE* in the sequenced *Escherichia coli* isolates. Amino acid substitutions are numbered relative to the corresponding protein of *E. coli* MG1655. A dash indicates that no substitution was detected in that gene. Asterisks mark substitutions identified here that have not previously been reported in association with fluoroquinolone resistance; the same substitution is marked wherever it occurs. Isolates EC47, EC112 and EC119 were used for the ciprofloxacin exposure experiments shown in Figures 5 and 6.

| **Isolate** | ***gyrA*** | ***gyrB*** | ***parC*** | ***pare*** |
|---|---|---|---|---|
| EC32 | S83L, D87N | – | S80I | S458A |
| EC47 | S83L, D87N | – | S80I | S458A |
| EC59 | – | – | E62K | – |
| EC60 | – | – | E62K | – |
| EC84 | D678E* | E185D* | D475E* | V136I* |
| EC87 | – | – | E62K | – |
| EC102 | – | – | E62K | – |
| EC112 | S83L, D87N | – | S80I | – |
| EC117 | S83L, D87N | – | S80I, E62K | S458A |
| EC119 | S83L, D87N | – | S80I, E62K | S458A |
| EC122 | S83L, D87N | – | E62K, S80I, E84V, A192V, A471G, Q481H, D475E* | – |
| EC127 | D678E*, A828S | E185D* | E62K, D475E* | V176I |
| EC141 | S83L, D87N | – | S80I, E62K, E84G | S458A |

Supplementation with 10 µg/mL ciprofloxacin, exposure well below the MIC of ≥ 128 µg/mL for all three isolates, markedly suppressed visually observed rdar morphotype expression in all three isolates (Figure 5A). At the same concentration, planktonic growth of *E. coli* EC112 in LB at 37 °C was indistinguishable from that of the untreated control (Figure 6D), indicating that the loss of the morphotype is not simply a consequence of growth inhibition; growth under the colony-assay conditions themselves (salt-free LB agar at 28 °C) was not measured. Ciprofloxacin treatment concomitantly abolished the characteristic web-like extracellular matrix architecture observed in rdar biofilms (Figure 5B). Under ciprofloxacin exposure, rdar-positive isolates exhibited a pink, dry and rough colony morphology (Figure 5A), which may reflect reduced curli production; curli and cellulose were, however, not quantified directly. Consistently, ciprofloxacin significantly reduced biofilm formation at both 28 °C and 37 °C (Figure 5C). Collectively, these findings suggest that ciprofloxacin exerts off-target secondary effects beyond its primary targets (GyrA, GyrB, ParC, and ParE), influencing rdar biofilm formation in *E. coli*.

**Figure 5.**
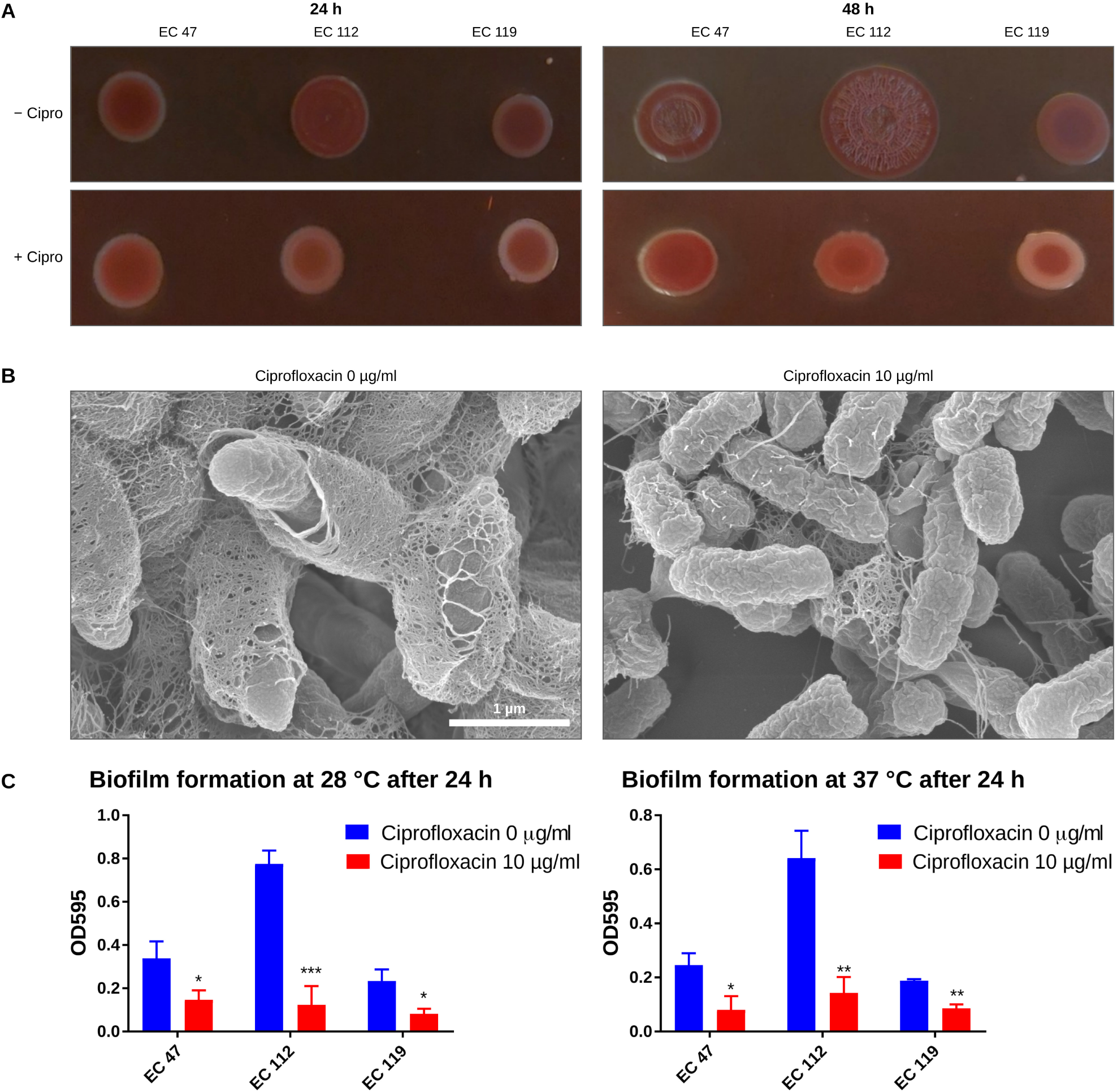
Ciprofloxacin suppresses rdar biofilm formation in ciprofloxacin-resistant *Escherichia coli*. (A) Representative images of rdar biofilm formation by ciprofloxacin-resistant *E. coli* isolates after 24 h (left) and 48 h (right) of growth on Congo red agar at 28 °C, in the absence (− Cipro) or presence (+ Cipro) of 10 µg/mL ciprofloxacin. (B) Scanning electron microscopy images of *E. coli* EC112 grown on Congo red agar for 48 h at 28 °C, illustrating disruption of rdar biofilm matrix architecture upon ciprofloxacin exposure. Scale bar, 1 µm, applies to both panels. (C) Quantification of biofilm formation by *E. coli* isolates grown in 96-well microtiter plates for 24 h at 28 °C and 37 °C, with and without 10 µg/mL ciprofloxacin. Error bars show mean ± SD of three independent experiments. Statistical significance was assessed by a two-tailed unpaired t-test with Welch’s correction: * *p* < 0.05, ** *p* < 0.01, *** *p* < 0.001.

### Ciprofloxacin inhibits colonization of ciprofloxacin-resistant *E. coli* on glass surfaces

Previously, we identified mountain-like biofilm patches formed by Csu pili and poly-N-acetylglucosamine (PNAG) produced by *Acinetobacter baumannii* on glass surfaces [33]. Because *Escherichia coli* does not harbor Csu pili, we sought to determine whether *E. coli* forms similar biofilm architectures on glass. To address this, *E. coli* EC112 was grown on glass surfaces, and live-cell phase-contrast microscopy was performed after 3 and 6 days of incubation at 28 °C.

Unlike *A. baumannii*, *E. coli* did not form discrete mountain-like biofilm patches. Instead, *E. coli* cells colonized the glass surface uniformly as a thin layer of adherent cells, covering approximately 50% of the surface after 3 days and approximately 80% after 6 days of incubation (Figure 6A). Consistent with our finding demonstrating that ciprofloxacin inhibits biofilm formation, supplementation of the growth medium with 10 µg/mL ciprofloxacin completely abolished surface colonization by *E. coli* EC112, despite its ciprofloxacin-resistant phenotype (Figures 6B, 6C).

**Figure 6.**
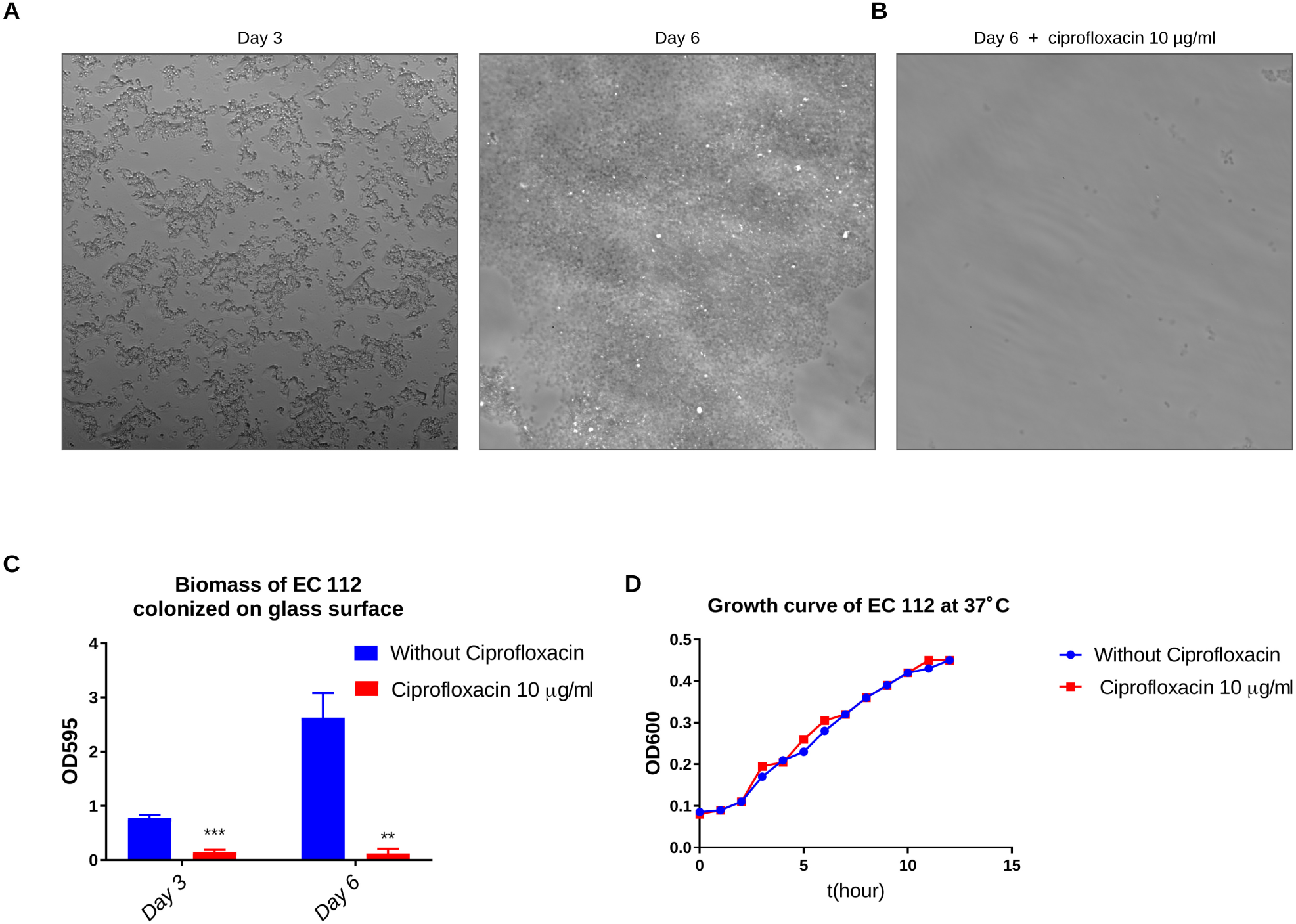
Ciprofloxacin inhibits surface colonization of ciprofloxacin-resistant *Escherichia coli*. (A) Representative phase-contrast live-cell images of *E. coli* EC112 colonizing glass surfaces after 3 and 6 days of incubation at 28 °C in salt-free LB medium. (B) Live-cell imaging of *E. coli* EC112 after 6 days of incubation on glass surfaces at 28 °C in salt-free LB medium in the presence of 10 µg/mL ciprofloxacin, showing inhibition of surface colonization. (C) Bar chart of biofilm mass of *E. coli* EC112 colonized on glass surface in the presence and absence of 10 µg/mL ciprofloxacin in salt-free LB medium. Error bars show mean ± SD. Statistical significance was assessed by a two-tailed unpaired t-test with Welch’s correction: ** *p* < 0.01, *** *p* < 0.001. (D) Growth curves of *E. coli* EC112 in LB medium with and without 10 µg/mL ciprofloxacin at 37 °C using the built-in temperature control mode of the Spark multimode plate reader (Tecan). The Y axis represents the optical density (OD600) measured at 60-min intervals for 12 hours. The experiment was done in triplicate and the curves were drawn using the mean OD600 values.

Altogether, these findings indicate that ciprofloxacin exerts a strong antibiofilm activity that suppresses surface colonization even in a ciprofloxacin-resistant isolate, highlighting a distinct inhibitory effect on early biofilm establishment on abiotic surfaces.

## Discussion

The red, dry and rough (rdar) morphotype represents a highly structured biofilm lifestyle defined by the coordinated production of the extracellular polysaccharide cellulose and amyloid curli fimbriae. Although well characterized in laboratory and environmental *E. coli* strains, its prevalence, regulation, and functional impact in clinical uropathogenic *E. coli* remain poorly understood. Here, we systematically characterize rdar morphotype expression in a large collection of clinical *E. coli* isolates and reveal important associations with antimicrobial susceptibility and fluoroquinolone-mediated modulation of biofilm architecture.

The prevalence of rdar morphotype expression in clinical *E. coli* has been debated. Early studies, often with limited sample sizes, reported sporadic rdar expression among clinical isolates [5, 38]. By analyzing a large, consecutively collected set of clinical *E. coli* isolates, we provide robust evidence that rdar-like biofilm traits are expressed by a substantial fraction of clinical strains. Although rdar expression was more frequently observed at 28 °C, a significant subset of isolates displayed rdar-like phenotypes at 37 °C. This observation suggests that, for many UPEC isolates, rdar expression is not strictly environmentally suppressed at host-relevant temperatures but instead is modulated through strain-specific regulatory networks and local cues, consistent with recent work on biofilm regulation in pathogenic *E. coli* [19].

One important and somewhat unexpected finding is the differential association between rdar morphotype expression and antimicrobial susceptibility. Although biofilm formation is often linked with increased tolerance to antibiotics, rdar-positive isolates were significantly more susceptible than smooth and white isolates to aztreonam, cefuroxime and cephradine. The relationship is not uniform, however: resistance to nitrofurantoin was significantly more frequent among rdar-positive isolates, and for most agents the two groups did not differ. The picture is therefore agent-specific rather than a simple trade-off, and it underscores that biofilm-forming capacity and planktonic antibiotic resistance are distinct phenotypes that need not be co-selected. Similar dissociations between biofilm formation and drug resistance have been observed in UPEC and other Gram-negative pathogens, where resistance is driven primarily by target mutations, efflux systems, or plasmid-borne determinants rather than biofilm-associated traits alone [39, 40].

Nitrofurantoin is a first-line oral agent for uncomplicated urinary tract infection in most treatment guidelines, and this result therefore deserves particular attention. Nitrofurantoin resistance was significantly more frequent among rdar-positive isolates (45.2% versus 20.2%, *p* = 0.0097) and among the cellulose-only pdar isolates (80.0% versus 20.2%, *p* = 0.010), but not among the curli-only bdar isolates, which points to an association with cellulose production rather than with curli, although the pdar group is small. If cellulose-producing isolates are indeed less often susceptible to nitrofurantoin, empirical use of this agent could select for such uropathogens. Confirmation of this hypothesis would require susceptibility data linked to treatment outcome, which our study did not investigate.

A central and clinically relevant observation of our study is the modulation of the rdar morphotype by ciprofloxacin. Exposure to ciprofloxacin suppressed rdar biofilm formation at a concentration that did not affect planktonic growth. These findings indicate that ciprofloxacin exerts off-target effects affecting biofilm-associated regulatory pathways beyond its canonical inhibition of DNA gyrase and topoisomerase IV.

Antibiotic-mediated modulation of biofilm regulatory networks has gained increasing attention [41, 42]. Subinhibitory concentrations of fluoroquinolones have been shown to alter global transcriptional programs, stress responses, and cyclic-di-GMP signaling in *E. coli* and other Gram-negative species [16, 43]. The shift from a red towards a pink colony morphology under ciprofloxacin exposure may indicate that curli production is more sensitive than cellulose production, but this remains to be confirmed by direct quantification of the matrix components. Such modulation may represent a bacterial adaptive response that remodels cell surface architecture under antibiotic stress, with potential consequences for surface attachment and community formation. Similar antibiofilm potential has been shown for clarithromycin in *Salmonella typhimurium* ATCC 14028 [44].

Whole-genome sequencing and phylogenetic analysis further showed that rdar morphotype expression spans diverse phylogroups and sequence types, with no clear lineage restriction. This reinforces the concept that rdar biofilm formation represents a conserved and flexible trait within the *E. coli* species rather than a feature confined to specific pathogenic clones. Recent comparative genomic studies also highlight extensive plasticity in biofilm regulatory circuits among UPEC strains [19, 20, 45, 46]; this may explain why even closely related isolates differ in rdar biofilm-related gene expression and regulatory responsiveness.

Sequence analysis of the *csgD* promoter region revealed polymorphisms distributed differently between isolates showing temperature-dependent and semi-constitutive rdar expression. Many of these polymorphisms map to predicted binding sites for global regulators such as IHF, OmpR, and CsgD itself. Within the CsgD/OmpR binding region the rdar-forming isolates of this collection share a sequence pattern largely distinct from that of the smooth and white isolates. We present these results as an association rather than a mechanism: no individual polymorphism segregates with phenotype once the previously characterized reference strains are included, and the polymorphisms do not track *csgD* transcript level. Fine-scale variation in regulatory sequences is nevertheless capable of producing major phenotypic differences in *Enterobacteriaceae* [47, 48]. Establishing which regulatory mechanisms are relevant for a specific isolate will require promoter-reporter fusions in an isogenic background and direct assessment of binding of the specific transcriptional regulators.

This study has several limitations. The isolates come from a single diagnostic laboratory in one city over a three-year period, so the prevalence figures reported here may not generalize to other settings or patient populations. No data reporting the clinical outcome were available, so the associations between rdar morphotype and susceptibility cannot be connected to treatment response. The polymorphisms identified in the *csgD*–*csgBA* intergenic region are associative: they were defined on a set of 23 sequences, no single position segregates with phenotype once previously characterized reference strains are included, and they were not tested in an isogenic background. Ciprofloxacin was tested only in ciprofloxacin-resistant isolates; whether a comparable antibiofilm effect occurs in susceptible isolates, at concentrations below their much lower MICs, was not examined. The effect of ciprofloxacin on rdar expression was examined in three isolates and on one strain for the surface-colonization and growth experiments, so the generality of the effect across the collection remains to be established. Some isolates in the collection share identical antimicrobial susceptibility and virulence profiles. These were recovered from different patients and are treated here as independent isolates. The occurrence of epidemiologically related isolates cannot, however, be excluded. Any such relatedness would reduce the effective number of independent observations underlying the between-morphotype comparisons. Finally, Congo red morphotypes were scored on solid medium and susceptibility was determined on planktonic cells, two conditions that need not report on the same physiological state.

## Conclusions

Our study establishes the rdar morphotype as a clinically relevant biofilm phenotype in clinical *E. coli* isolated from urine and shows that subinhibitory ciprofloxacin suppresses rdar biofilm formation in ciprofloxacin-resistant isolates. These findings highlight the need to consider antibiotic-induced alterations in bacterial multicellular behavior when interpreting antimicrobial susceptibility and treatment outcomes. Future work should elucidate the molecular mechanisms underlying ciprofloxacin-mediated suppression of the rdar morphotype and determine whether similar regulatory effects occur in vivo during UTI therapy. A deeper understanding of how antibiotics reshape biofilm regulatory networks may inform the development of improved strategies to prevent and treat persistent and recurrent urinary tract infections.

## Supporting information

upplementary_Information.pdf

## Abbreviations

bdar: brown, dry and rough
c-di-GMP: cyclic di-GMP
CFU: colony-forming units
CLSI: Clinical and Laboratory Standards Institute
ESBL: extended-spectrum β-lactamase
IHF: integration host factor
LB: lysogeny broth
MIC: minimum inhibitory concentration
MLST: multilocus sequence typing
OD: optical density
PBS: phosphate-buffered saline
pdar: pink, dry and rough
PNAG: poly-N-acetylglucosamine
qRT-PCR: quantitative reverse transcription PCR
SD: standard deviation
Rdar: red, dry and rough
Saw: smooth and white
SEM: scanning electron microscopy
ST: sequence type
UPEC: uropathogenic *Escherichia coli*
UTI: urinary tract infection

## Declarations

### Ethics approval and consent to participate

The study was approved by the institutional ethical review committee of the University of Health Sciences, Lahore (letters UHS/Education/126-16/ and UHS/REG-18/ERC). Verbal informed consent was obtained from all participants, and the committee waived written documentation of patient information because the isolates and their associated genotypic and phenotypic data cannot be traced to individual patients.

### Consent for publication

Not applicable.

### Availability of data and materials

The whole-genome sequence data generated in this study have been deposited in the NCBI Sequence Read Archive under BioProject accession PRJNA1554061. All other data supporting the findings of this study are included in this article and its additional files.

### Authors’ contributions

IAh conceived and designed the study, acquired funding, performed data analysis and wrote the manuscript. FM and IAl performed the experiments, analyzed the data and wrote the original draft. HS, RY and AB performed experiments. BEU and UR contributed to study design, supervision and funding acquisition. All authors read, revised and approved the final manuscript. Throughout this section, IAh denotes Irfan Ahmad and IAl denotes Irfan Ali.

## Acknowledgements

Not applicable.

## Additional files

**Additional file 1: Table S1. Sequence types and distribution of antimicrobial resistance and virulence genes among the whole-genome-sequenced isolates.**

Sequence types, antimicrobial resistance genes and virulence genes identified by whole-genome sequencing in the fourteen clinical isolates selected to span the range of Congo red morphotypes.

**Additional file 2: Table S2. Distribution of antimicrobial resistance genes in relation to Congo red morphotype.**

Comparative distribution of the antimicrobial resistance genes detected by whole-genome sequencing, grouped by Congo red morphotype.

**Additional file 3: Table S3. Distribution of virulence genes in relation to Congo red morphotype.**

Comparative distribution of the virulence genes detected by whole-genome sequencing, grouped by Congo red morphotype.

Additional file 4: Figure S1. Unmodified scanning electron micrograph of wild-type *Escherichia coli* No. 12

The wild-type micrograph of Figure 1B as acquired, at its native magnification (14.06 nm per pixel) and without the cropping and resampling applied in Figure 1B. Scale bar, 1 µm.

**Additional file 5: Figure S2. Complete multiple sequence alignment of the *csgD*–*csgBA* intergenic region**

Full ClustalW alignment of the *csgD*–*csgBA* intergenic region for the 23 sequences analyzed in Figure 4, comprising the 14 whole-genome-sequenced study isolates and 9 previously characterized reference strains retrieved from the database. Positions are numbered relative to the translation start site of *csgD* in *E. coli* MG1655. Isolates are ordered by phenotypic class and polymorphic positions are shaded; the regions shown in Figure 4A–D are boxed.

**Additional file 6: Figure S3. Virulence determinants and extended-spectrum β-lactamase production in relation to Congo red morphotype**

Prevalence of the virulence determinants *afa*, *fimH*, *pap*, *sfa*, K1 and K5, and of extended-spectrum β-lactamase (ESBL) production, among clinical *E. coli* isolates grouped by Congo red morphotype (*n* = 146). Bars show the percentage of tested isolates that were positive and the number of positive isolates is given above each bar. ESBL is a phenotype rather than a gene and is separated from the virulence determinants on the axis.

Additional file 7: Table S4. Previously described *Escherichia coli* strains used in this study.

Reference strains whose *csgD*–*csgBA* intergenic sequences were included in the alignment shown in Figure 4, with their phylogroup, rdar phenotype and source.

