## Supplementary material for "Regulatory plasticity of the rdar biofilm morphotype in clinical uropathogenic *Escherichia coli* and its modulation by ciprofloxacin": upplementary_Information.pdf

**Contents**

Additional file 1: Table S1. Sequence types and distribution of antimicrobial resistance and virulence genes among the whole-genome-sequenced isolates

Additional file 2: Table S2. Distribution of antimicrobial resistance genes in relation to Congo red morphotype

Additional file 3: Table S3. Distribution of virulence genes in relation to Congo red morphotype

Additional file 4: Figure S1. Unmodified scanning electron micrograph of wild-type Escherichia coli No. 12

Additional file 5: Figure S2. Complete multiple sequence alignment of the csgD–csgBA intergenic region

Additional file 6: Figure S3. Virulence determinants and extended-spectrum β-lactamase production in relation to Congo red morphotype

Additional file 7: Table S4. Previously described Escherichia coli strains used in this study

**Additional file 4: Figure S1. Unmodified scanning electron micrograph of wild-type Escherichia coli No. 12**

Figure S1. E. coli No. 12 wild type, native magnification (14.06 nm per pixel)

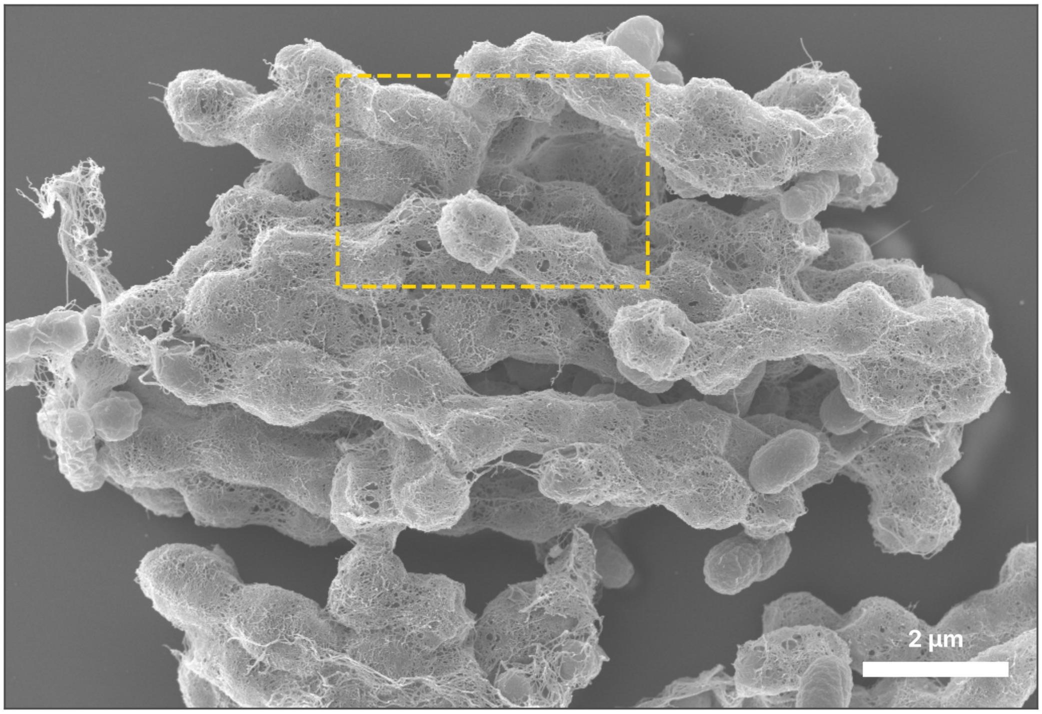

The wild-type micrograph of Figure 1B as acquired, at its native magnification (14.06 nm per pixel) and without the cropping and resampling applied in Figure 1B. Scale bar, 1 µm.

**Additional file 5: Figure S2. Complete multiple sequence alignment of the csgD–csgBA intergenic region**

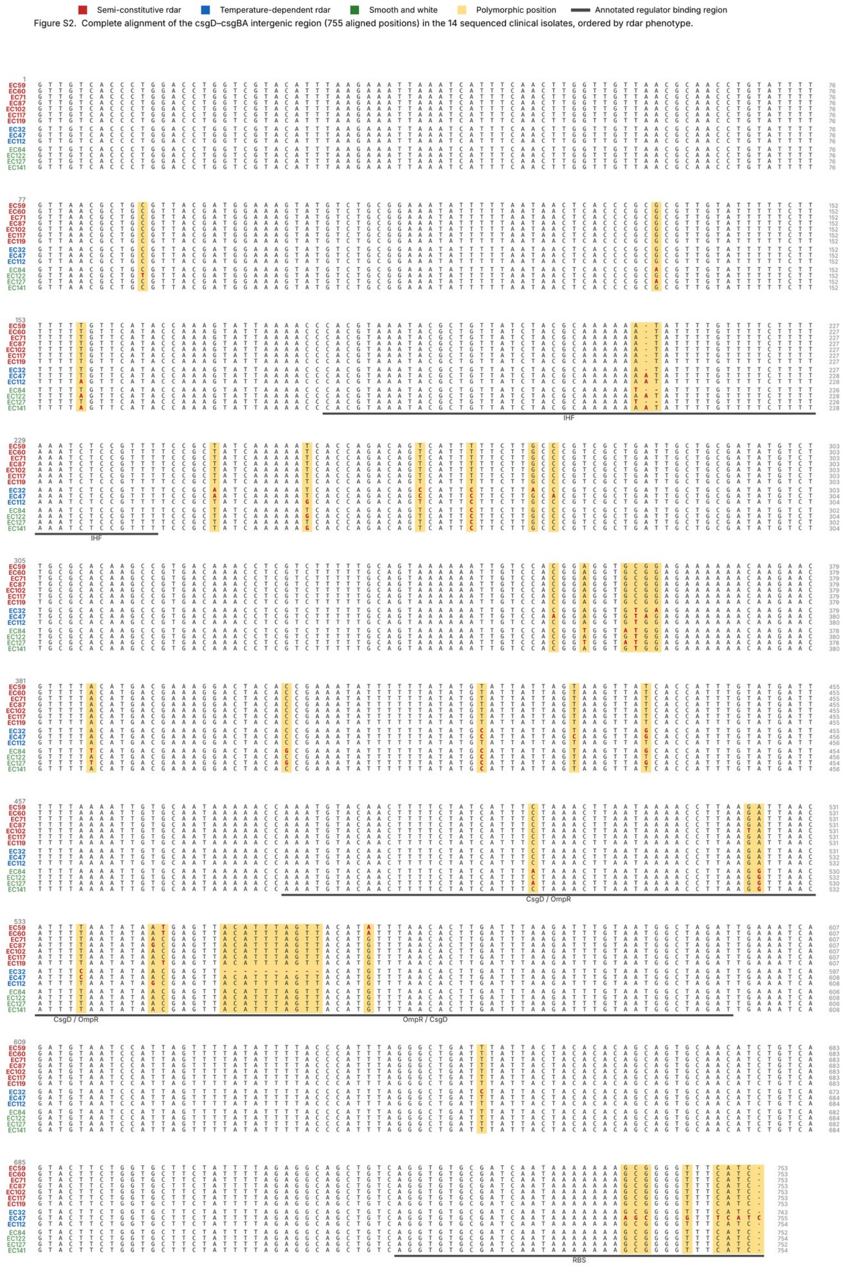

Full ClustalW alignment of the csgD–csgBA intergenic region for the 23 sequences analyzed in Figure 4, comprising the 14 whole-genome-sequenced study isolates and 9 previously characterized reference strains retrieved from the database. Positions are numbered relative to the translation start site of csgD in E. coli MG1655. Isolates are ordered by phenotypic class and polymorphic positions are shaded; the regions shown in Figure 4A–D are boxed.

**Additional file 6: Figure S3. Virulence determinants and extended-spectrum β-lactamase production in relation to Congo red morphotype**

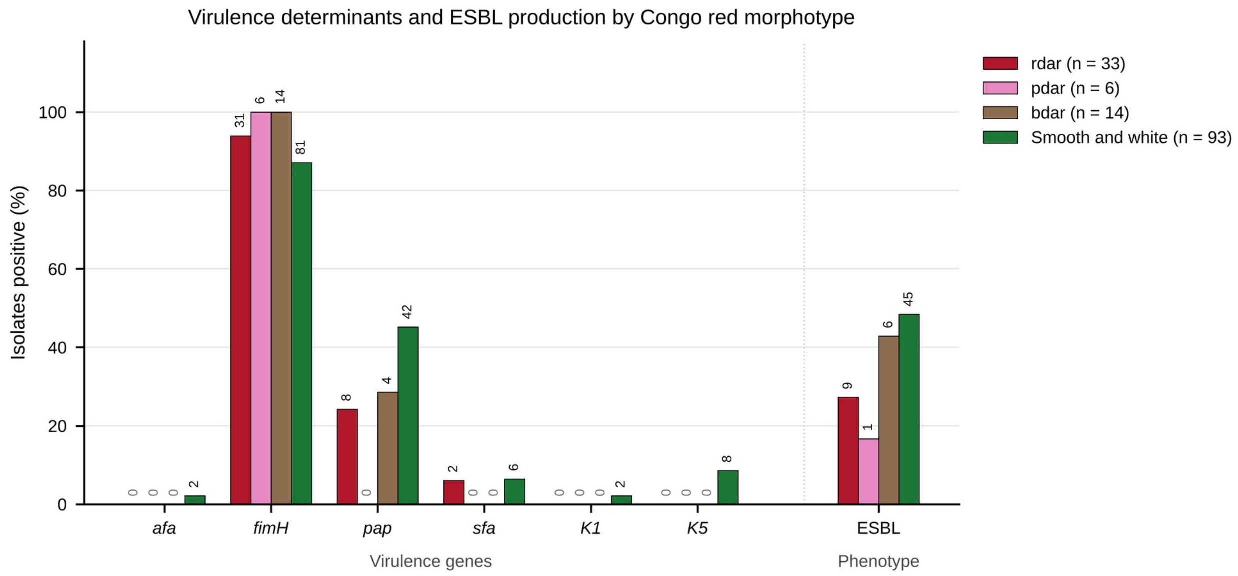

Prevalence of the virulence determinants afa, fimH, pap, sfa, K1 and K5, and of extended-spectrum β-lactamase (ESBL) production, among clinical E. coli isolates grouped by Congo red morphotype (n = 146). Bars show the percentage of tested isolates that were positive and the number of positive isolates is given above each bar. ESBL is a phenotype rather than a gene and is separated from the virulence determinants on the axis.

**Additional file 1: Table S1. Sequence types and distribution of antimicrobial resistance and virulence genes among the whole-genome-sequenced isolates**

| | **MLST Result**<br>**Seq type** | **Allele** | **Drug resistant features** | **Drug Resistant genes** | **Virulence factors** |
|---|---|---|---|---|---|
| EC 71 | 58 | *adk_6*<br>*fumC_4*<br>*gyrB_4*<br>*icd_16*<br>*mdh_24*<br>*purA_8*<br>*recA_14* | Fosfomycin | *fosA7* | *capU, gad, hlyE, iss*<br>*kpsE*<br>*kpsMII_K5*<br>*lpfA*<br>*ompT*<br>*terC* |
| | | | Quinolone | *qnrS1* | |
| | | | Folate pathway antagonist | *dfrA14*<br>*sul2* | |
| | | | Tetracycline | *tet(A)* | |
| | | | Beta-lactam | *blaTEM-1B* | |

| | | | | | |
|---|---|---|---|---|---|
| EC87 | 12548 | *adk_9*<br>*fumC_19*<br>*gyrB_22*<br>*icd_26*<br>*mdh_11*<br>*purA_8*<br>*recA_2* | | | *gad*<br>*hlyE*<br>*lpfA*<br>*terC* |
| EC 102 | 2175 | *adk_6*<br>*fumC_4*<br>*gyrB_15*<br>*icd_88*<br>*mdh_43*<br>*purA_8*<br>*recA_14* | | | *astA*<br>*cba*<br>*cma*<br>*etsC*<br>*gad*<br>*hlyE*<br>*lpfA*<br>*ompT*<br>*terC*<br>*traT* |
| EC 112 | 12548 | *adk_939**<br>*fumC_19**<br>*gyrB_22* | Aminoglycoside | *aadA1*<br>*aph(6)-Id*<br>*aph(3'')-Ib* | *fyuA*<br>*gad*<br>*hlyE*<br>*irp2*<br>*lpfA*<br>*terC* |
| | | | Aminocyclitol | *aadA1*<br>*aadA1* | |
| | | | Quinolone | *gyrA:p.D87N*<br>*gyrA:p.S83L* | |
| | | | Folate pathway antagonist | *sul2* | |
| | | | Tetracycline | *tet(B)* | |
| | | | Beta-lactam | *blaTEM-1B*<br>*blaOXA-1* | |
| | | | Amphenicol | *catA1* | |
| EC59 | 1727 | *adk_6*<br>*fumC_19*<br>*gyrB_3*<br>*icd_16*<br>*mdh_11*<br>*purA_8*<br>*recA_6* | Fosfomycin | *fosA7* | *capU*<br>*gad*<br>*hlyE*<br>*lpfA*<br>*terC* |
| | | | Aminoglycoside | *aph(6)-Id*<br>*aph(3'')-Ib* | |
| | | | Quinolone | *qnrS1* | |
| | | | Folate pathway antagonist | *sul2*<br>*dfrA14* | |
| | | | Tetracycline | *tet(A)* | |
| | | | Beta-lactam | *blaTEM-1B*<br>*blaCTX-M-15* | |
| EC 32 | 405 | *adk_35*<br>*fumC_37*<br>*gyrB_29*<br>*icd_25*<br>*mdh_4*<br>*purA_5* | Folate pathway antagonist | *sul1*<br>*dfrA17* | *air*<br>*chuA*<br>*eilA*<br>*fyuA*<br>*gad*<br>*hlyE* |
| | | | Aminoglycoside | *aph(6)-Id*<br>*aph(3'')-Ib*<br>*aac(6')-Ib-cr*<br>*aac(3)-IIa* | |

| | | | | | |
|---|---|---|---|---|---|
| | | recA_73 | | aadA5<br>aph(6)-Id | irp2<br>iss<br>iutA<br>kpsE<br>kpsMII_K5<br>ompT<br>papA_F43<br>sitA<br>terC<br>traT<br>usp<br>yfcV |
| | | | Macrolide | mph(A) | |
| | | | Quinolone | aac(6')-Ib-cr | |
| | | | Aminocyclitol | aadA5 | |
| | | | Tetracycline | tet(B) | |
| | | | Beta-lactam | blaOXA-1<br>blaCTX-M-15 | |
| | | | Quaternary ammonium compound | QacE | |
| | | | Amphenicol | catB3<br>catB3 | |
| EC 117 | 90 | adk_6<br>fumC_4<br>gyrB_12<br>icd_1<br>mdh_20<br>purA_8<br>recA_7 | Folate pathway antagonist | sul2<br>dfrA17<br>sul1 | hlyE<br>iss<br>senB<br>terC<br>traT |
| | | | Aminoglycoside | aph(3'')-Ib<br>aac(6')-Ib-cr<br>aph(6)-Id<br>aadA5 | |
| | | | Tetracycline | tet(A) | |
| | | | Macrolide | mph(A) | |
| | | | Quinolone | aac(6')-Ib-cr | |
| | | | Aminocyclitol | aadA5 | |
| | | | Rifamycin | ARR-3 | |
| | | | Beta-lactam | blaOXA-1<br>blaCTX-M-27 | |
| | | | Quaternary ammonium compound | QacE | |
| | | | Amphenicol | catB3 | |
| EC 47 | 648 | adk_92<br>fumC_4<br>gyrB_87<br>gyrB_87<br>mdh_70<br><br>purA_58<br>recA_2 | Streptogramin b | erm(B | afaD<br>air<br>chuA<br>fyuA<br>gad<br>hlyE<br>irp2<br>iss<br>iucC<br>iutA<br>kpsE<br>kpsMII<br>lpfA<br>ompT<br>sitA<br>terC<br>traT<br>yfcV |
| | | | Aminoglycoside | aadA5 | |
| | | | Folate pathway antagonist | dfrA17<br>sul1 | |
| | | | Quinolone | parE:p.S458A<br>gyrA:p.S83L<br>gyrA:p.D87N<br>parC:p.S80I | |
| | | | Aminocyclitol | aadA5 | |
| | | | Tetracycline | tet(B) | |
| | | | Beta-lactam | blaTEM-1B<br>blaOXA-1<br>blaCMY-42<br>blaCTX-M-15 | |
| | | | Macrolide | mph(A)<br>erm(B) | |
| | | | Quaternary ammonium compound | QacE | |
| | | | Amphenicol | catB3 | |

| | | | | | |
|---|---|---|---|---|---|
| | | | | *catB3* | |
| | | | Lincosamide | *erm(B)* | |
| EC 119 | 424 | *adk_6*<br>*fumC_30*<br>*gyrB_32*<br>*icd_16*<br>*mdh_11*<br>*purA_8*<br>*recA_7* | Macrolide | *mph(A)* | *cib*<br>*gad*<br>*hlyE*<br>*lpfA*<br>*terC*<br>*ter T* |
| | | | Aminoglycoside | *aph(3'')-Ib*<br>*aph(6)-Id*<br>*aadA2* | |
| | | | Quinolone | *qnrS1*<br>*qepA4* | |
| | | | Folate pathway antagonist | *sul1*<br>*dfrA12*<br>*sul2* | |
| | | | Tetracycline | *tet(A)* | |
| | | | Aminocyclitol | *aadA2* | |
| | | | Beta-lactam | *blaOXA-181*<br>*blaTEM-1B* | |
| | | | Quaternary ammonium compound | *QacE* | |
| EC 141 | 167 | *adk_10*<br>*fumC_11*<br>*gyrB_4*<br>*icd_8*<br>*mdh_8*<br>*recA_2*<br>*purA_13* | Aminoglycoside | *aac(6')-Ib-cr*<br>*aadA5* | *capU*<br>*gad*<br>*hlyE*<br>*iss*<br>*terC*<br>*traT* |
| | | | Aminocyclitol | *aadA5* | |
| | | | Quinolone | *aac(6')-Ib-cr* | |
| | | | Folate pathway antagonist | *sul1*<br>*dfrA17* | |
| | | | Tetracycline | *tet(A)* | |
| | | | Beta-lactam | *blaCTX-M-15*<br>*blaOXA-1*<br>*blaCMY-42* | |
| | | | Macrolide | *mph(A)* | |
| | | | Quaternary ammonium compound | *QacE* | |
| | | | Amphenicol | *catB3*<br>*catB3* | |
| EC 53 | 131 | *adk_53*<br>*fumC_40*<br>*gyrB_47*<br>*icd_13*<br>*mdh_36*<br>*purA_28*<br>*recA_29* | Aminoglycoside | *aac(6')-Ib-cr* | *cea, chuA, fyuA, gad, hlyA, iha, irp2, Iis, iucC, iutA, kpsE, kpsMII_K5, ompT, papA_F43, sat, sitA, terC, usp, yfcV, cnf1,* |
| | | | Quinolone | *aac(6')-Ib-cr* | |
| | | | Peroxide | *SitABCD* | |
| | | | Beta-lactam | *blaOXA-1* | |
| | | | Amphenicol | *catB3*<br>*catB3* | |
| EC60 | 1727 | *adk_6*<br>*fumC_19*<br>*gyrB_3*<br>*icd_16*<br>*mdh_11*<br>*purA_8*<br>*recA_6* | Fosfomycin | *fosA7* | *capU, gad2, hlyE, lpfA, terC* |
| | | | Aminoglycoside | *aph(6)-Id*<br>*aph(3'')-Ib* | |
| | | | Quinolone | *qnrS1* | |
| | | | Folate pathway antagonist | *sul2*<br>*dfrA14* | |
| | | | Tetracycline | *tet(A)* | |
| | | | Beta-lactam | *blaTEM-1B* | |

| | | | | | |
|---|---|---|---|---|---|
| | | | | *blaCTX-M-15* | |
| EC 84 | 73 | *adk_36*<br>*fumC_24*<br>*gyrB_9*<br>*icd_13*<br>*purA_11*<br>*mdh_17*<br>*recA_25* | Peroxide | *SitABCD* | *astA*, *cea*,, *chuA*, *clbB*, *cnf1*, *focC*, *safe*,<br>*focG*, *foci*, *fyuA*, *fyuA*, *hra*, *iroN*, *irp2*,<br>*iss*, *kpsE*, *kpsMII_K23*, *mchB*, *mchC*, *mchF*, *mcmA*,<br>*ompT*, *pic*, *sfaD*, *sitA*,<br>*tcpC*, *usp*, *vat*, *yfcV* |
| EC 104 | 1823 | *adk_64*<br>*fumC_7*<br>*gyrB_5*<br>*icd_83*<br>*mdh_8*<br>*recA_6*<br>*purA_8* | | | *cma*, *gad*, *hlyE*, *iss*<br>*lpfA*<br>*ompT*<br>*papC*<br>*terC*<br>*traT* |
| EC 122 | 10 | *adk_10*<br>*fumC_11*<br>*gyrB_4*<br>*icd_8*<br>*mdh_8*<br>*purA_8*<br>*recA_2* | Aminoglycoside | *aadA2* | *cea*, *fyuA*, *gad*, *hlyE*,<br>*hra*, *iha*, *irp2*, *iss*, *iutA*, *kpsE*, *kpsMII_K1*, *mchB*,<br>*mchC*, *mchF*, *neuC*,<br>*papA_feiA_F8*, *sat*, *senB*,<br>*sitA*, *terC*, *traT* |
| | | | Macrolide | *mph(A)* | |
| | | | Quinolone | *qepA4* | |
| | | | Folate pathway antagonist | *sul1*<br>*dfrA12* | |
| | | | Peroxide | *SitABCD* | |
| | | | Aminocyclitol | *aadA2* | |
| | | | Beta-lactam | *blaCTX-M-15* | |
| | | | Quaternary ammonium compound | *QacE* | |
| EC 127 | 73 | *adk_36*<br>*fumC_24*<br>*gyrB_9*<br>*icd_13*<br>*mdh_17*<br>*purA_11*<br>*recA_25* | Aminoglycoside | *aph(6)-Id*<br>*aph(3'')-Ib* | *cea*, *chuA*, *clbB*, *cnf1*<br>*Foc*, *csfaE*, *focG*, *focI*<br>*fyuA*, *gad*, *hlyA*, *hra*<br>*Iha*, *ireA*, *iroN*, *irp2*<br>*Iss*, *iucC*, *iutA*, *kpsE*<br>*kpsMII_K5*, *mchB*<br>*mchC*, *mchF*, *mcmA*<br>*ompT*, *papA_F43*<br>*papA_F9*, *papC*, *pic*<br>*sat*, *sitA*, *tcpC*, *usp*<br>*vat*, *yfcV* |

**Additional file 2: Table S2. Distribution of antimicrobial resistance genes in relation to Congo red morphotype**

| **Drug resistant genes** | **Drug** | **Rdar**<br>**N=7**<br><br>**MLST:**<br>**58, 12548, 2175, 12548, 1727, 405, 90** | **Saw**<br>**N=7**<br><br>**MLST**<br>**10, 1823, 73, 131, 167, 424, 648** |
|---|---|---|---|

| | | | |
|---|---|---|---|
| *fosA7* | Fosfomycin | 3 | 0 |
| *qnrS1* | Quinolone | 3 | 1 |
| *dfrA14*<br>*sul2* | Folate pathway antagonist<br>Sulfonamide | 3<br>4 | 1<br>3 |
| *tet(A)* | Tetracycline | 3 | 4 |
| *aadA1*<br>*aph(6)-Id*<br>*aph(3'')-Ib* | Aminoglycoside | 3<br>3<br>3 | 0<br>5<br>4 |
| *gyrA:p.D87N*<br>*gyrA:p.S83L* | Quinolone | 1<br>1 | 1<br>1 |
| *Sul1* | Sulfonamide | 0 | 6 |
| *tet(B)* | Tetracycline | 1 | 2 |
| *blaTEM-1B*<br>*blaOXA-1* | Beta lactam<br>Beta lactam | 4<br><br>1 | 3<br>5 |
| *dfrA17* | Folate pathway antagonist | 0 | 4 |
| *aac(6')-Ib-cr* | Quinolone | 0 | 6 |
| *aac(3)-IIa* | Aminoglycosides | 0 | 6 |
| *aadA5* | Aminoglycosides | 0 | 6 |
| *mph(A)* | Macrolide | 0 | 6 |
| *tet(B) 2* | Tetracycline | 0 | 2 |
| *blaCTX-M-15* | Beta lactam | 0 | 5 |
| *QacE* | Quaternary ammonium compound | 0 | 6 |
| *catB3* | Amphenicol | 0 | 6 |
| *ARR-3* | Rifamycin | 0 | 1 |
| *blaCTX-M-27* | Aminoglycosides | 0 | 1 |
| *qnrB1* | Quinolone | 0 | 1 |
| *SitABCD* | Peroxide | 0 | 2 |
| *aadA2* | Aminocyclito | 0 | 4 |
| *blaOXA-181* | Beta-lactam | 0 | 1 |
| *blaCMY-42* | Aminocyclitol | 0 | 2 |
| *erm(B)* | Lincosamide | 0 | 1 |

**Additional file 3: Table S3. Distribution of virulence genes in relation to Congo red morphotype**

| **VIRULENCE FACTOR** | **PROTEIN FUNCTION** | **Rdar**<br>**N=7**<br><br>**MLST:**<br>**58, 12548, 2175, 12548, 1727, 405, 90** | **Saw**<br>**N=7**<br><br>**MLST**<br>**10, 1823, 73, 131, 167, 424, 648,** |
|---|---|---|---|
| *cea* | Colicin E1 | 0 | 5 |
| *chuA* | Outer membrane hemin receptor | 1 | 5 |
| *clbB* | Hybrid non-ribosomal peptide / polyketide megasynthase | 0 | 2 |
| *cnf1* | Cytotoxic necrotizing factor | 0 | 3 |
| *focCsfaE* | | 0 | 2 |
| *focG* | F1C adhesion | 0 | 2 |

| | | | |
|---|---|---|---|
| *Foci* | S fimbrial/F1C minor subunit | 0 | 2 |
| *fyuA* | Siderophore receptor | 2 | 6 |
| *hlyA* | Haemolysin A | 0 | 2 |
| *Hra* | Heat-resistant agglutinin | 0 | 3 |
| *Iha* | Adherence protein | 0 | 3 |
| *ireA* | Siderophore receptor | 0 | 1 |
| *iroN* | Enterobactin siderophore receptor protein | 0 | 2 |
| *irp2* | High molecular weight protein 2 non-ribosomal peptide synthetase | 2 | 6 |
| *Iss* | Increased serum survival | 2 | 7 |
| *iucC* | Aerobactin synthetase | 0 | 4 |
| *iutA* | Ferric aerobactin receptor | 0 | 5 |
| *kpsE* | Capsule polysaccharide export inner-membrane protein | 2 | 6 |
| *kpsMII_K5* | Polysialic acid transport protein; Group 2 capsule | 2 | 6 |
| *mchB* | Microcin H47 part of colicin H | 0 | 3 |
| *mchC* | MchC protein | 0 | 3 |
| *mchF* | ABC transporter protein MchF | 0 | 3 |
| *mcmA* | Microcin M part of colicin H | 0 | 2 |
| *ompT* | Outer membrane protease (protein protease 7) | 4 | 6 |
| *papA_F43* | Major pilin subunit F43 | 1 | 3 |
| *papA_F9* | Major pilin subunit F9 | | |
| *papC* | Outer membrane usher P fimbriae | 0 | 2 |
| *Pic* | serine protease autotransporters of Enterobacteriaceae (SPATE) | 0 | 2 |
| *Sat* | Secreted autotransporter toxin | 0 | 3 |
| *sitA* | Iron transport protein | 1 | 6 |
| *tcpC* | Tir domain-containing protein | 0 | 2 |
| *Usp* | Uropathogenic specific protein | 1 | 3 |
| *Vat* | Vacuolating autotransporter toxin | 0 | 2 |

### Additional file 7: Table S4. Previously described Escherichia coli strains used in this study

▶ AUTHOR CHECK: references for the strains marked '[ref required]' must be supplied before submission.

| Strain | Description | Phylogroup | rdar phenotype | Source / reference |
|---|---|---|---|---|
| E. coli No. 12 (UPEC12) | Uropathogenic isolate; well-characterised rdar morphotype former; parent of the ΔcsgA, ΔbcsA and ΔbcsA ΔcsgA derivatives used in Figure 1 | B2 | rdar | Kai-Larsen et al. 2010 [24]; biofilm maturation characterised by Zhang et al. 2024 [48] |
| E. coli No. 12 ΔcsgA | Curli-deficient derivative of E. coli No. 12 | B2 | pdar (cellulose only) | This laboratory [ref required] |
| E. coli No. 12 ΔbcsA | Cellulose-deficient derivative of E. coli No. 12 | B2 | bdar (curli only) | This laboratory [ref required] |
| E. coli No. 12 ΔbcsA ΔcsgA | Cellulose- and curli-deficient derivative | B2 | saw | This laboratory [ref required] |
| E. coli MG1655 | K-12 laboratory reference strain; reference genome for REALPHY and for numbering of csgD promoter positions | A | saw | Laboratory collection [ref required] |
| E. coli EcoR31 | ECOR reference collection; semi-constitutive rdar morphotype | — | semi-constitutive rdar | Cimdins et al. 2017 [20]; Cimdins-Ahne et al. 2023 [21] |
| E. coli B11870 | Commensal/ uropathogenic isolate; semi-constitutive rdar morphotype | B2 | semi-constitutive rdar | Cimdins et al. 2017 [20]; Cimdins-Ahne et al. 2023 [21] |
| E. coli Fec101 | Phylogroup B1 reference strain | B1 | rdar | [ref required] |
| E. coli Fec67 | Reference strain | — | rdar | [ref required] |
| E. coli TOB1 | Phylogroup B2 reference strain | B2 | rdar | [ref required] |
| E. coli B8638 | Phylogroup D reference strain | D | rdar | [ref required] |
